# Identification of molecular determinants of gene-specific bursting patterns by high-throughput imaging screens

**DOI:** 10.1101/2024.06.08.597999

**Authors:** Varun Sood, Ronald Holewinski, Thorkell Andresson, Daniel R. Larson, Tom Misteli

## Abstract

Stochastic transcriptional bursting is a universal property of active genes. While different genes exhibit distinct bursting patterns, the molecular mechanisms for gene-specific stochastic bursting are largely unknown. We have developed and applied a high-throughput-imaging based screening strategy to identify cellular factors and molecular mechanisms that determine the bursting behavior of human genes. Focusing on epigenetic regulators, we find that protein acetylation is a strong acute modulator of burst frequency, burst size and heterogeneity of bursting. Acetylation globally affects the Off-time of genes but has gene-specific effects on the On-time. Yet, these effects are not strongly linked to promoter acetylation, which do not correlate with bursting properties, and forced promoter acetylation has variable effects on bursting. Instead, we demonstrate acetylation of the Integrator complex as a key determinant of gene bursting. Specifically, we find that elevated Integrator acetylation decreases bursting frequency. Taken together our results suggest a prominent role of non-histone proteins in determining gene bursting properties, and they identify histone-independent acetylation of a transcription cofactor as an allosteric modulator of bursting via a far-downstream bursting checkpoint.

## Introduction

Transcription is highly stochastic^1–4^. Rather than exhibiting constant activity, RNA polymerases engage at the gene promoter stochastically to generate temporary bursts of RNA (On-time), but transcriptional activity intermittently ceases for periods of time (Off-time)^5–9^. This phenomenon of stochastic transcriptional bursting is wide-spread, evolutionarily conserved^1, 2, 4, 10, 11^ and represents a major source of intrinsic transcriptional noise amongst individual cells in a population^9^. The regulation of stochastic bursting has been implicated in critical functions such as cell fate, development and responses to environmental stimuli and diseases^12–14^.

Bursting patters are gene specific and highly varied^11, 15, 16^. Native genes can differ in both the On- and/or Off-times often with larger variations observed for the Off-times^16^. The mean On-times typically vary between 10-20 min while the mean Off times can range between 20 min to 100 min for a typical fast and slow bursting gene, respectively, and can extend to days for certain genes^16, 17^. Bursting patters can be altered, for example by gene induction or suppression, however, it is unclear how changes in On- and Off-times in response to a stimulus differ between genes within a cell or for the same gene across different cell types and how these changes impact the overall expression levels of the affected genes^11, 18–21^.

Several modulators of transcription bursting have been identified, including interaction of transcription factors and *cis*-regulatory elements at both proximal^22^ and distal^23^ sites, as well as the cellular levels of trans-activators^24, 25^, *cis*- element architecture^11, 26^, chromatin accessibility^27, 28^ and DNA supercoiling^29, 30^ have all been linked directly or indirectly to gene bursting.

Epigenetic modifications also affect bursting. The basic unit of chromatin, the nucleosome, decreases bursting by restricting access to transcriptional machinery^28, 31^ and decreasing transcription factor dwell times on chromatin^22, 32^. Epigenetic modifications are thought to act by modulating nucleosome stability and/or RNA pol II pause release^33^, both of which can impact bursting^28, 31, 34^. Indeed, different epigenetic marks correlate with^35^ or, in some cases, modulate bursting kinetics^36^. For example, methylation of histone 3 at lysine 4 in the 5’ region of *act5* and *sdc* is required for both inheritance and maintenance of burst frequency in *Dictyostelium*^37, 38^, and increased global histone acetylation increases burst size of multiple genes in several cell lines^11, 18, 19^. Furthermore, guided chromatin modifications support a direct role in bursting: dCas9-directed acetylation of the *BMAL1* promoter in NIH3T3 cells and *FOS* enhancer in N2A cells increases burst frequency^20^ and burst size^21^, respectively. Thus, chromatin modifications can act as a dynamic rheostat for gene-specific bursting patterns^36^. On the other hand, genome-wide comparison of active marks like acetylation of histone 3 on lysine 27 at the gene promoters show a relatively weak correlation with bursting^39^ as bursting of several genes remains unchanged or even decreases with an increase in histone acetylation^11, 40^. In addition, epigenetic modifications such as acetylation and methylation often also target non-histone proteins and while several mechanisms have been proposed for how these protein modifications affect transcription, their impact on bursting kinetics are unknown^41–48^.

The identification of cellular mechanisms that control bursting is impeded by major experimental challenges. The most accurate measurements of bursting dynamics are obtained from live-cell gene reporters often using fluorescently labelled RNAs as a read out^6^. However, these approaches rely on engineered cell lines which limits the number of genes that can be analyzed. Genome-wide approaches, such as single-cell RNA-seq, on the other hand are not suitable for genes with low expression nor can they be used in unbiased screening approaches to identify new regulatory factors^49^. To overcome some of these limitations and to comprehensively identify epigenetic mechanisms that directly impact bursting we have developed a high-throughput RNA-FISH based imaging assay to measure bursting of endogenous genes at single-cell resolution. We used this assay in a screen to assess the effect of a wide spectrum of epigenetic modifications on bursting of fourteen genes in multiple human cell lines testing thousands of experimental perturbations. We find that protein acetylation is a strong acute modulator of bursting with differential impact on the On- and Off-times. However, the effects of acetylation are not mediated via histone modifications since promoter histone acetylation was not correlated with bursting changes. Rather, we show that acetylation of subunit 4 of the Integrator complex plays a causal role in bursting. These results demonstrate a major role of chromatin-independent acetylation on gene bursting, and they point to a far downstream checkpoint in control of gene stochasticity.

## Results

### Imaging-based single-cell high-throughput screen for effectors of gene bursting

We sought to comprehensively examine the direct impact of epigenetic modifications on bursting of multiple endogenous genes across different cell lines by high-throughput screening. This approach necessitates fast perturbation of chromatin states coupled with an assay to quantitatively detect bursting behavior at the single-allele level. To this end, and to be able to probe endogenous genes, we developed a high-throughput compatible, sensitive and fast fluorescence in situ hybridization (FISH) assay (Fig. 1). The assay is based on detection of nascent transcripts by RNA-FISH (nscRNA-FISH) with probes targeting introns in the genes of interest to capture active transcription sites (TS; Fig. 1A) and uses the frequency of TS in the cell population as a first approximation for the burst behavior of the gene of interest. For example, the number of detectable TS in a population of cells will be higher for a gene with short Off-times and long On-times as opposed to gene with long Off-times and short On-times (Fig. 1A). Moreover, this amplification-free technique enables quantitative interpretation of TS intensity. As proof of principle, we directly compared FISH detection with an established MS2-GFP based live-cell assay using the well characterized *ERRFI1* gene in the diploid transformed human bronchial epithelial cells (HBECs)^16^, where treatment with the deacetylase inhibitor trichostatin A (TSA) resulted in a marked increase in overall bursting (Fig. S1)^50^. These changes were reflected in an increase in both the number and intensity of visible MS2-GFP TS as early as 2 h and fully evident at 4 h of treatment (Fig. 1B). A similar increase in TS frequency and intensity at *ERRFI1* was detected using nscRNA-FISH (Fig. 1C). The fold increase in TS frequency between control and TSA treated sample was marginally larger in the live-cell assay compared to the FISH assay whereas the fold increase in intensity measurements were similar in the two assays (Fig. 1B and 1C). We conclude that nscRNA-FISH faithfully detects changes in gene bursting behavior.

**Figure 1:**
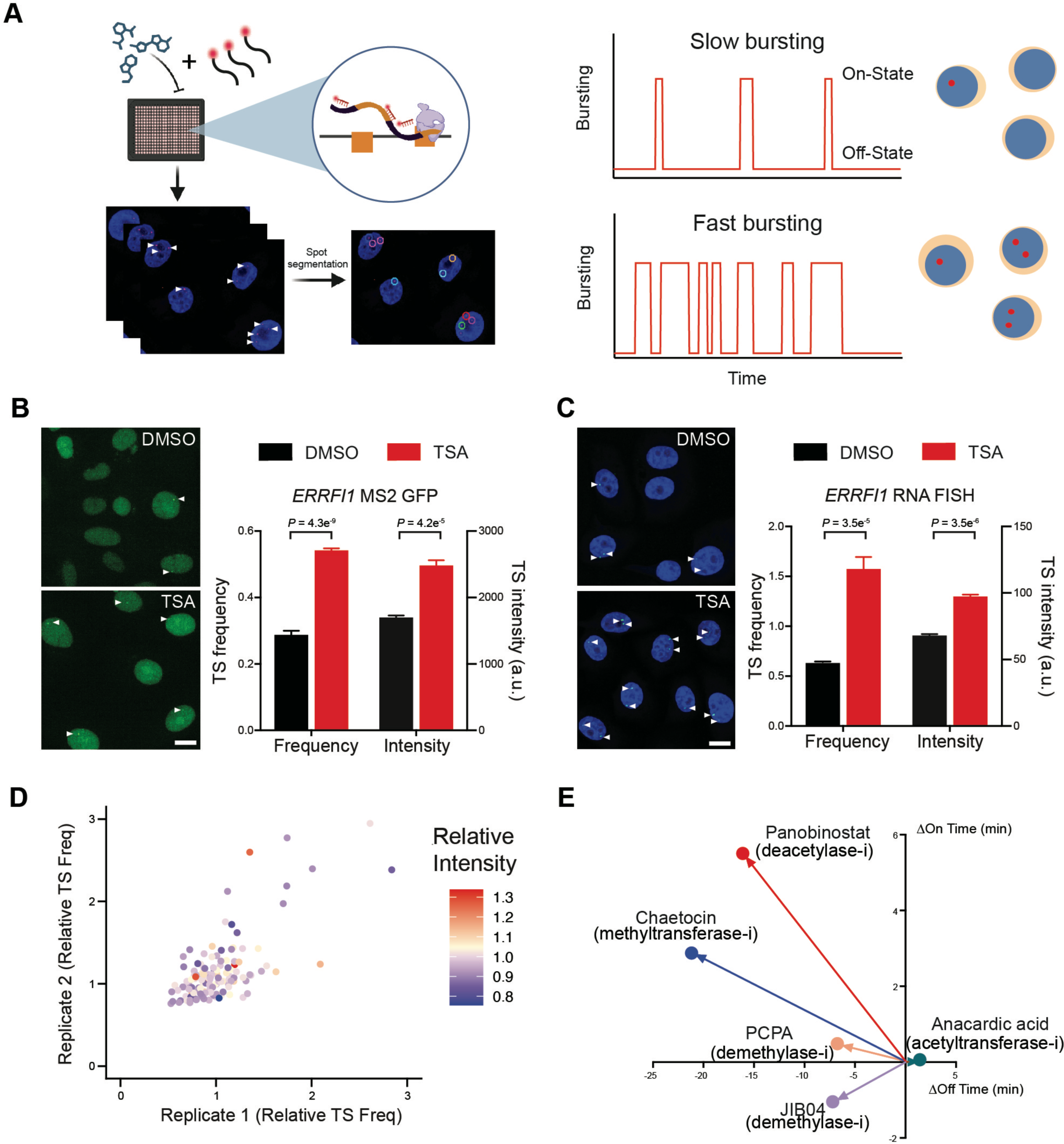
A high-throughput assay for transcriptional bursting of native genes based on nscRNA-FISH. **A.** Schematic of a high-throughput nscRNA-FISH assay to measure transcriptional bursting of endogenous genes. Left. Fluorescent probes targeting the intron of target genes are hybridized to fixed cells in 384-well plate and the number of active TS are measured using automated image acquisition and spot segmentation. Right. The average number of active TS for cells in the population is a surrogate for the bursting behavior of genes. The number of active TS at any given time in the population is higher for a gene with high bursting rate. **B** and **C**. The RNA-FISH assay accurately detects the increase in transcription bursting seen in live cells. Representative image (left) and plot (right) for the average frequency and intensity of *ERRFI1* TS detected using either MS2-GFP (B) or nscRNA-FISH (C) in HBEC for control (DMSO) or trichostatin A (TSA) treated cells. Values represent mean ± SEM of >300 cells from 3 biological replicates. Scale bar = 10μm and *P* value: two sample t-test. Active transcription sites in **A**, **B** and **C** are indicated by arrowheads. **D.** Reproducibility of small molecule modulator screen for *ERRFI1* bursting. Relative TS frequency (Freq) of *ERRFI1*, defined as the ratio of TS frequency in treated vs untreated (DMSO) controls from two biological replicates of the screen. The average change in the relative TS intensity is color coded on each data point. **E**. Changes in the mean Off- and On-times from live-cell bursting assay for *ERRFI1* recoded between 4 h to 16 h of treatment with selected inhibitors. Data is representative of >30 cells from one or more acquisitions.

As proof-of-principle for its suitability in a screen, we next used the FISH-based bursting assay in an automated high-throughput screen to assess the effect of over a hundred perturbations on the bursting behavior of *ERRFI1*. We first assayed bursting changes for *ERRFI1* in response to a library of 140 small molecule epigenetic modulators that were predominantly inhibitors, in a 384-well format using automated fluidic handling for cell seeding, transfer of media and solutions, and use of automated imaging and image analysis pipeline (Fig. 1A; Table S1; see Methods). The small molecule library targets twenty classes of epigenetic modifiers with at least two modulators for ten classes and multiple modulators representing the major classes of epigenetic modifiers including acetyltransferases, deacetylases, methyltransferases, demethylases, bromodomain proteins, sirtuins and DNA methyltransferases (DNMT) (see Fig. S2 for details). We intentionally used a small molecule library, rather than RNAi or CRISPRi perturbation, and limited treatment to 4 h to assess acute effects of perturbations on gene bursting rather than long-term epigenetic changes and to reduce the probability of secondary effects. A total of 43/140 (∼31%) and 32/140 (∼23%) treatments led to significant changes in *ERRFI1* TS frequency and intensity, respectively, based on a 95% confidence limit of changes in Z-scores (see Methods for details; Table S2) with good agreement between screen replicates (Fig. 1D).

To validate primary hits based on nscRNA-FISH, we selected five representative inhibitors that altered *ERRFI1* TS frequency and assayed real-time bursting kinetics using an established MS2-GFP based live-cell *ERRFI1* bursting assay^51^. HBECs with monoallelic insertion of the viral MS2 operator array in *ERRFI1* intron^16^ were incubated with the selected inhibitors for 4 h and bursting was assayed for the next 12 h (Fig. 1E). As expected, depending on the inhibitor, we see different combinations of changes in Off- and On-time of *ERRFI1* bursting. For example, the deacetylase inhibitor panobinostat increased *ERRFI1* bursting by decreasing Off-time and increasing On-time, whereas the demethylase inhibitor JIB04 decreased both Off- and On-time (Fig. 1E). Despite their different timescales of action, we observe good correspondence between the short-term bursting changes observed in the nscRNA-FISH screen and long-term bursting changes in the live-cell assay (Fig. S3). For example, an increase in bursting of *ERRFI1* with panobinostat observed in both assays is consistent with a decrease in Off-time and increase in On-time confirming the nscRNA-FISH results. We conclude that FISH-based detection of TS frequency faithfully reports the acute response to epigenetic modulators in a high-throughput format.

### Identification of effectors of bursting dynamics

To get insights into general features of bursting dynamics we repeated the bursting screen in a set of 14 genes with a wide range of known bursting rates and gene expression states^16, 52^. The gene set includes both high and low expressors (Fig. 2A) and real-time bursting data is available in human bronchial epithelial cells (HBECs) for nine genes^16^ (*ERRFI1, CANX, DNAJC5, RPAP3, SEC16A, SLC2A1, KPNB1, RAB7A* and *RHOA*), whereas five additional genes were selected based on their known clinical or physiological relevance (*EGFR, MYC, NRAS, P53* and *GAPDH*), but without available live-cell bursting data. We also extended the screen to two additional epithelial cell lines, the breast cancer cell line MCF7 and the human colon cancer cell line HCT116, that differ in tissue of origin, gene expression profiles, and chromatin landscape^52^. Using the small molecule library of 140 compounds and probing 14 genes in each cell line, a total of 5,880 conditions were tested in the screen and the change in TS frequency relative to control measured after 4 h of treatment (Fig. 2A). Out of the 1960 conditions tested in each cell line, significant changes (*P* <0.05) were observed in 414 (21%), 361 (18%) and 643 (33%) conditions in HBECs, MCF7 and HCT116, respectively, with the probability of bursting changes being highly gene specific (Table S3). Comparison between cell lines revealed that a major fraction of significant changes were cell-type specific and showed little covariation; significant changes in TS frequency for the same gene by the same modulator in two or three cell lines represented <25% or 8% of hits, respectively (Fig S4A and S4B). Taken together, these results show that bursting of genes within the same cell differed in sensitivity to epigenetic modifications and showed cell-specific changes.

**Figure 2.**
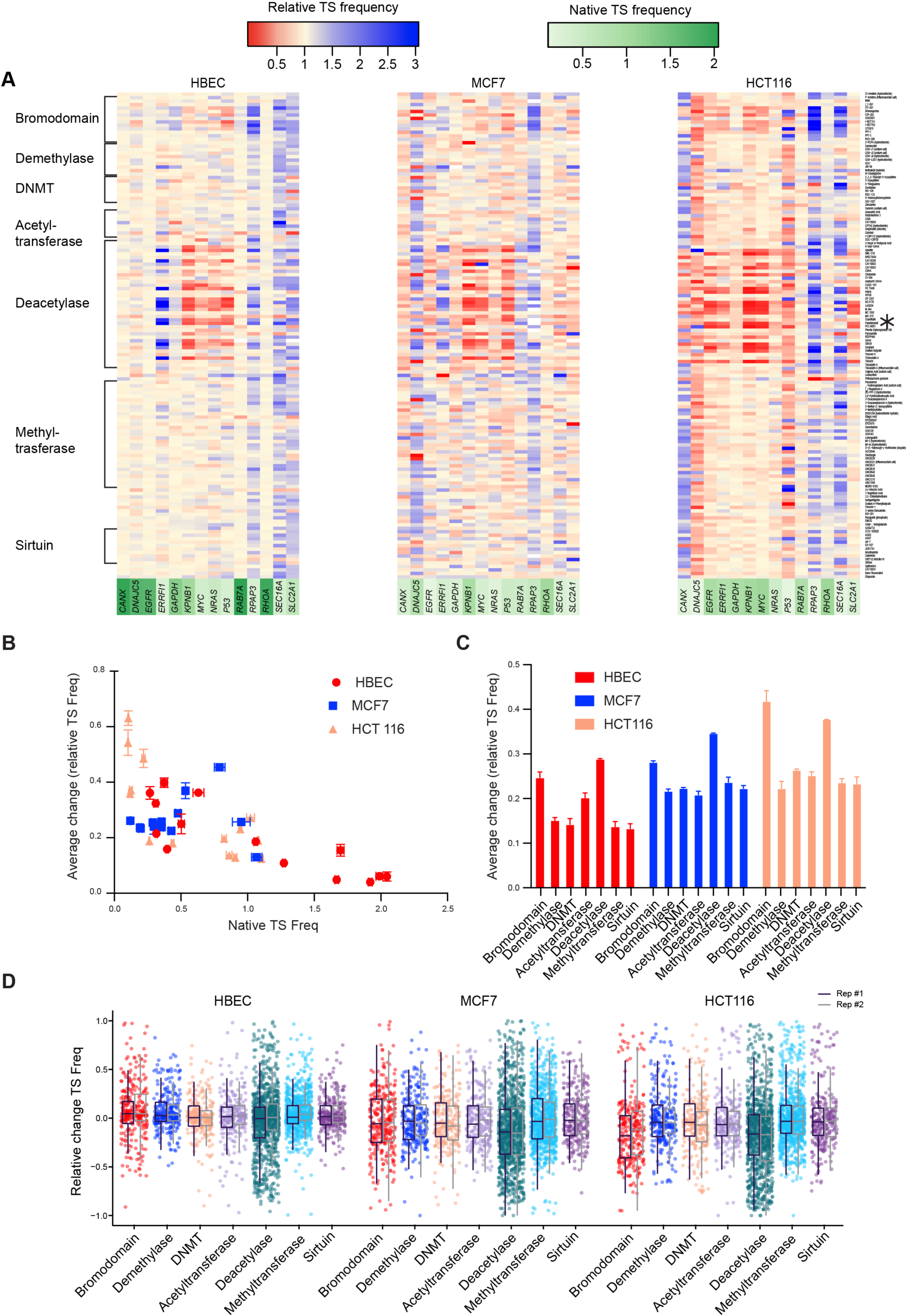
TS frequency changes are sensitive to changes in acetylation and inversely related with native TS frequencies across multiple cell lines. **A**. Heat maps of the relative TS frequency (Freq), defined as the ratio of TS frequency in treated vs untreated (DMSO) controls (Freq_Treated_/Freq_Untreated_) obtained from the screen with small molecule modulators targeting different classes of chromatin modifying enzymes for model genes in three cell types: HBEC, MCF7 and HCT116. Treatments with no detectable TS are colored white. The rows are arranged by modulators of major classes indicated on the left, panobinostat treatment is asterisked on the right and gene names are color coded based on their native TS frequencies. Data represents mean of two biological replicates. **B**. Relationship between native TS frequency (Native TS Freq) and the average change in relative TS frequency, defined as the magnitude of TS frequency changes relative to the native TS frequency for a gene averaged across the screen, avg(|Freq_Treated_-Freq_Untreated_|)/Freq_Untreated_. Values represent mean ± SEM of at least ten and two biological replicates for Freq_Untreated_ and the average magnitude change, respectively. **C** and **D**. Quantitation of the impact of the major classes of chromatin modifying enzymes on TS frequency changes. **C**. Comparison of the magnitude of change in relative TS frequency, averaged across all modulators in the respective classes, avg(|Freq_Treated_-Freq_Untreated_|)/Freq_Untreated_, in three cell lines. Values represent mean ± SEM of two biological replicates. **D**. Relative changes in TS frequency, (Freq_Treated_-Freq_Untreated_)/Freq_Untreated_, for each modulator of the respective classes in the three cell lines. Values from the two biological replicates are shown adjacent. All TS frequency measurements were derived from imaging of >300 cells per condition and data shown for relative change between 1 and −1.

Strikingly, five genes (*RAB7A*, *RHOA*, *CANX*, *DNAJC5* and *EGFR*) in HBEC were largely insensitive to most modulators in the library (Fig. 2A, left). These genes shared the common property of being highly expressed based both on high TS frequencies (TS ∼2) in untreated fixed cells and frequent bursting in live cells (mean rate 0.028min^-1^)^16^, indicating that genes with high bursting rates might be less sensitive to changes in bursting upon epigenetic perturbation. To test this hypothesis more broadly, we compared the native TS frequency (number of active TS under control conditions) to the average change in relative TS frequency observed across the screen (Fig. 2B). The latter parameter was obtained by averaging the magnitude of relative changes from native TS frequency for each gene across all treatments. We observe a strong negative relationship between native TS frequency and sensitivity to modulators in HBEC and HCT116, and a slightly weaker relationship in MCF7 (Fig. 2B). In support, genes with larger changes in a principal component analysis (PCA) of relative TS frequency have on average lower native TS frequency compared to rest of the genes (0.33 vs 0.81; Fig. S4C). Our observation is in line with reports that strongly expressed genes are regulated by multiple pathways and tend to show little expression change in response to single perturbations^53, 54^. These findings point to a prominent role for native bursting levels in determining the sensitivity to epigenetic changes.

One of the key objectives of our screen was to uncover the impact of different chromatin modifications on bursting. To quantitatively address this question, we computed the changes in relative TS frequency for all genes by modulators of the seven major classes of epigenetic modifiers (bromodomain proteins, demethylases, DNA methyltransferases (DNMT), acetyltransferases, deacetylases, methyltransferases and sirtuins) in each cell line. Focusing on the extent of change, we averaged the magnitude of change in each class and found a greater impact on bursting by inhibitors of deacetylases, bromodomain proteins and acetyltransferases, in decreasing order of magnitude (Fig. 2C). The class-wise changes in the magnitude of bursting were generally higher in HCT116 and MCF7 cells compared to HBEC, which agrees with the higher average TS frequency for the model genes in HBEC (∼2-fold higher). Further, factoring in the directionality of the relative changes, we did not see a strong bias towards either an increase or decrease in bursting (Fig. 2D). Inspection of individual genes indicated that the directionality of change was both gene- and cell-line specific. For example, the TS frequency of *ERRFI1* was increased by deacetylase inhibitors like panobinostat, in HBEC and MCF7, but decreased in HCT116 (Fig. 2A). On the other hand, *MYC* TS frequency was generally decreased by panobinostat and other deacetylase inhibitors in all three cell lines (Fig. 2A). Given that these changes are visible as early as 4 h post treatment, they are likely a direct effect of the modulator induced changes. Taken together these findings identify acetylation as a strong global modulator of gene bursting.

### Effectors of transcription site intensity

The nscRNA-FISH signal intensity is directly proportional to the mRNA counts at the TS and is affected by burst size, burst frequency and gene length^34, 50, 55^. Thus, as an additional measure of bursting change, we determined the changes in TS intensities relative to the untreated samples (Fig. 3A). Like TS frequency changes, significant changes in TS intensity (p<0.05) were also both gene- and cell-type specific. Depending on the cell line, significant bursting changes occurred in a fraction of all treatments and varied between 12-36% (Fig. 3A and S5, Table S4). Furthermore, significant changes for the same gene with the same modulator between two or three cell lines showed a 14-22% or 4% overlap, respectively (Fig. S5).

**Figure 3.**
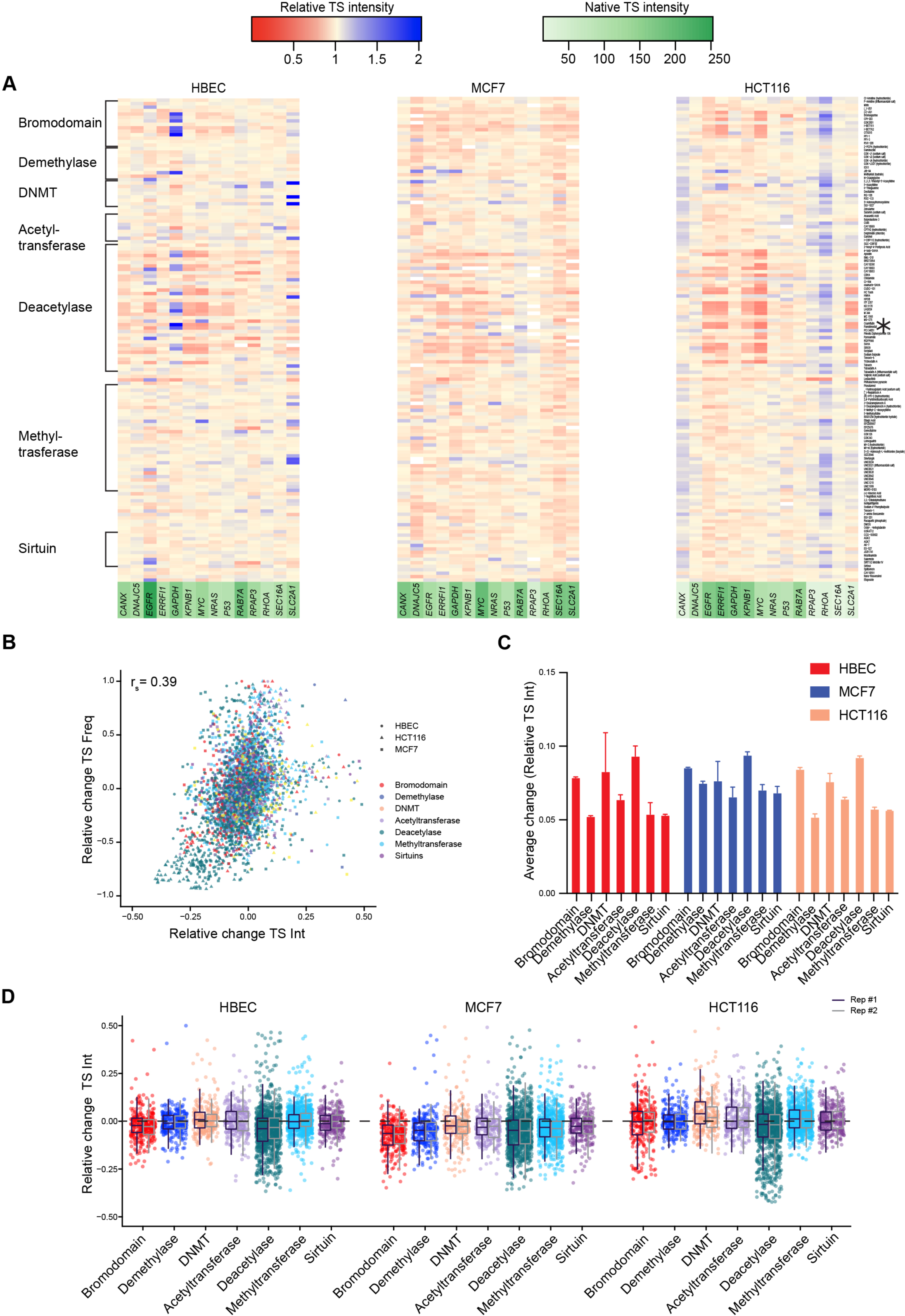
TS intensity changes are sensitive to acetylation and correlated with changes in TS frequency. **A**. Heat maps of the relative TS intensity (Int), defined as the ratio of TS intensity in treated vs untreated (DMSO) controls (Int_Treated_/Int_Untreated_) obtained from the screen with small molecule modulators targeting different classes of chromatin modifying enzymes for model genes in three cell types: HBEC, MCF7 and HCT116. Treatments with no detectable TS are colored white. The rows are arranged by modulators of major classes indicated on the left, panobinostat treatment is asterisked on the right and gene names are color coded based on their native TS intensities. Data represents mean of two biological replicates. **B**. Relationship between relative changes in TS frequency (Int_Treated_-Int_Untreated_)/Int_Untreated_) and TS intensity ((Int_Treated_-Int_Untreated_)/Int_Untreated_) in response to modulators of the major classes. Spearman rank coefficient (r_s_) indicated. Values represent mean of two biological replicates between ranges of −1.0 to 1.0 and −0.5 to 0.5 for frequency and intensity changes, respectively. **C** and **D**. Quantitation of the impact of the major classes of chromatin modifying enzymes on TS intensity changes. **C**. Comparison of the average change in relative TS intensity, defined as the magnitude of TS intensity changes relative to the native TS intensity averaged across all modulators in the respective classes, avg(|Int_Treated_- Int_Untreated_|)/Int_Untreated_, in three cell lines. Values represent mean ± SEM of two biological replicates. **D**. Relative changes in TS intensity, (Int_Treated_-Int_Untreated_)/Int_Untreated_, for each modulator in the respective classes from three cell lines. Values from the two biological replicates are show adjacent. All TS intensity measurements were derived from imaging of >300 cells per condition and data shown for relative change between 0.5 and −0.5.

Across the screen we find that the changes in the TS frequency and intensity, rather than their native values, were positively correlated (r_s_ =0.39) with 32%, 18% and 50% overlap between hits in HBEC, MCF7 and HCT116 cells, respectively (Fig 3B and S6A). Parsing these changes by major classes of modifier yielded similar levels of correlation while parsing by TS frequency yielded higher correlation for genes in the middle of the bursting spectrum (native frequency range 0.5-1.5) relative to genes on either end (Fig. S6B, S6C). The latter agrees with the expected increase in the number of nascent transcripts over the gene-body detected by nscRNA-FISH upon increase in burst frequency. Thus, TS frequency and intensity are more often regulated independently, however, instances of coregulation are influenced by the native bursting state of the gene rather than the class of modulator. Indeed, unlike TS frequency, we did not see any significant correlation between native TS intensity and the average magnitude of relative change in TS intensity across the screen for individual genes (Fig. S6D). As with TS frequency, however, we did find that inhibitors of deacetylases and bromodomain proteins, but not acetyltransferases, have the biggest impact on TS intensity (Fig. 3C) and generally lead to a decrease in TS intensity (Fig. 3D). Estimation of changes in heterogeneity for TS intensity (Fano factor=σ^2^/ µ) within the cell population (Fig. S7A) showed a strong decrease with deacetylase inhibitors (Fig. S7C and S7D) and a strong positive correlation between changes in heterogeneity and TS intensity (Fig. S7B, s = 0.74). Taken together, our results strongly point to a direct role of acetylation in regulating multiple aspects of transcriptional bursting.

### Dynamics of bursting and chromatin acetylation changes upon deacetylase inhibition

Having identified protein acetylation as the major modulator of bursting, we investigated the precise nature of the changes in bursting kinetics using live-cell imaging. We specifically asked how changes in protein acetylation affect On- and Off-times. For this analysis, we selected the clinically relevant pan-deacetylase inhibitor, panobinostat, since it was consistently identified in our screens amongst the most potent modulators of gene bursting (Fig. 2A and 3A). For live-cell observations, we used previously established HBEC lines with monoallelic insertions of MS2 operator arrays in the introns of *ERRFI1*, *KPNB1*, *RHOA*, *RAB7A* or *DNAJC5*^16^ since these genes covered a wide range of responses to panobinostat (Fig. 2A and 3A). In line with previous studies, we see a broad distribution of Off- and On-times for all genes ranging between ∼5-200 min and ∼2-50 min^15, 16^, respectively (Fig. 4A and 4B). Panobinostat consistently led to a decrease in the Off-time of all genes (Fig. 4A and 4C) in agreement with reported effects of increased protein acetylation on gene bursting^18, 20^. Upon closer inspection, we found that panobinostat mostly decreased the fraction of longer Off-times indicated by the disappearance of the shoulder population in the 75-120 min range (Fig. 4A) and an increase in the fraction of lower Off-times with a peak around 5-15 min. This change is unlikely due to a lack of sensitivity to detect smaller Off-time differences since similar pattern persisted at shorter acquisition intervals (Fig. S8A). Furthermore, this non-uniform change is also reflected by larger differences between the mean vs the median changes in Off-times, since the median is less sensitive to skewed distributions (Fig. S8B). We see a similar, but weaker, pattern for On-time changes (Fig. 4B, S8B) with a longer On-time shoulder around 20-30 min. Clusters of high, moderate, and low bursting genes have previously been reported to similarly differ in their total fraction of longer Off-times^16^ and we find that Off-time changes induced by representative inhibitors of acetyltransferases, methyltransferases and demethylases follow a similar pattern (Fig. S9). In summary, bursting changes induced by acetylation and other epigenetic changes preferentially act on the longer Off-times, and deacetylase inhibition uniformly increases the bursting frequency. Our data also suggests the existence of a lower biological limit or refractory period of around 5 min for the interval between two bursts.

**Figure 4.**
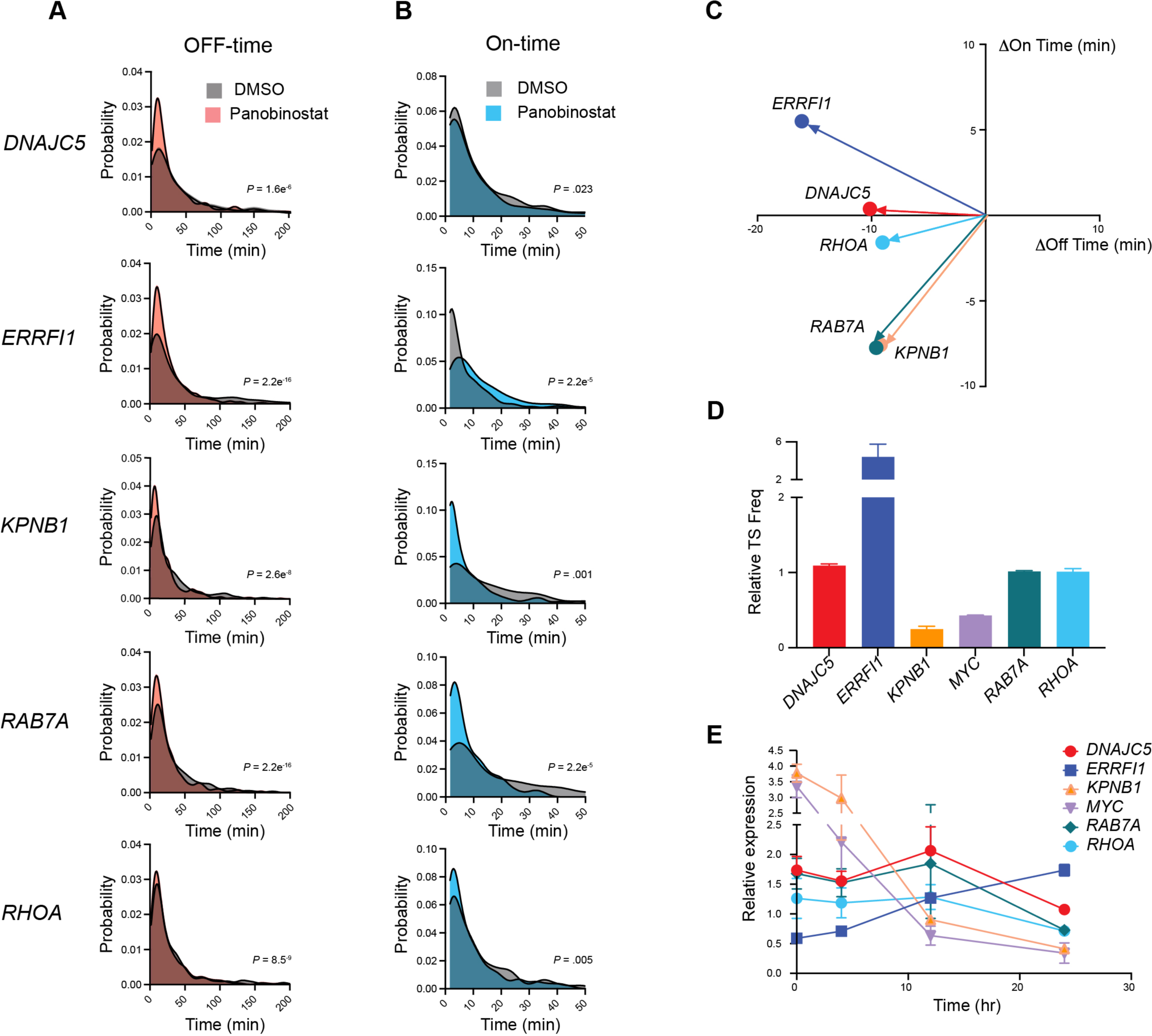
Deacetylase inhibitor decreases Off-time but change On- time on a gene-specific basis. **A** and **B**. Normalized histograms for the Off- (A) and On-(B) time of bursting for model genes in DMSO controls and deacetylase inhibitor (Panobinostat) treated cells from live-cell bursting assay using a monoclonal cell lines with MS2 inserted in the intron of model genes. The acquisition post 4 h of treatment was done for 12 h at 100 s interval. *P* value: Kolmogorov–Smirnov test. **C**. Mean changes in the Off- and On- time between panobinostat treated cell vs controls from live-cell bursting assay described in **A** and **B**. The data in **A-C** was derived from >40 cells from one or more acquisitions. **D**. Relative TS frequencies between panobinostat treated and control cells from nscRNA-FISH for model genes after four hours of incubation with panobinostat. Data represents mean of two biological replicates of >300 cells per condition **E**. qRT-qPCR for model genes at different incubations times in panobinostat. The abundances between samples were normalized with a spike-in control. Values represent mean ± SEM of at least three biological replicates.

In contrast to the Off-time changes, the On-time changes with panobinostat were more variable between genes. The absolute changes in On-times varied widely from +5 min (+57%) for *ERRFI1* to −7.6 min (−47%) for *KPNB1* with a mean of −2.4 ±2.06 min. In relative terms, the variations were even larger with a mean of −13.8 ± 16.2 % (Fig. 4C). On the other hand, the decrease in Off-times with respect to untreated controls varied from 9.6 min for *RAB7A* to 16 min for *ERRFI1* (Fig. 4C) with an average of 10.4 ±1.72 min. Expressed as percentage change these values were very similar across genes with a mean of 27.3 ±1.96 %. We thus asked if the On-time changes play a dominant role in determining the overall change in gene-specific bursting and expression levels. To test this, we assayed mRNA levels by spike-in normalized RT-qPCR for six model genes in the presence of panobinostat (Fig. 4E) and found that the On-time changes indeed correlated strongly with the mRNA levels after 24 h treatment. The expression was upregulated for *ERRFI1*, which was the only gene that exhibited an increase in On-time, whereas the other genes were downregulated with different kinetics that were related to their native bursting rates (Fig. 4D and E). For example, high-bursting genes like *RHOA*, *RAB7A* and *DNAJC5* were slow to repress, as they remained unchanged up until 12 h of treatment (Fig. 4E). The same genes also did not show any change in bursting after four hours in panobinostat (Fig. 2A and 4D). The comparatively slow busters, like *MYC* and *KPNB1*, on the other hand, showed early decreases in both bursting and expression levels after 4 h (Fig. 4D and 4E). In sum, increased protein acetylation leads to similar relative decreases in Off-times but diverse gene-specific changes in On-times with the latter playing a prominent role in determining overall mRNA levels.

### Promoter H3K27ac levels are stoichiometrically uncoupled from bursting changes

Based on our observations and the well-established role of acetylation of core histones in promoter nucleosomes^56–59^, we hypothesized that the observed modulation of gene bursting occurred via promoter-associated histone acetylation. To investigate the relationship between chromatin acetylation and bursting we assayed the changes in the ubiquitously associated histone mark for gene activation^52^, acetylation of histone 3 lysine 27 ac (H3K27ac), by spike-in calibrated ChIP-seq after 4 h of panobinostat treatment in HBECs. As expected, we saw a sharp increase in the nuclear levels of H3K27ac (Fig. 5A inset). At the metagene level this increase was localized to either the gene body or upstream of the −1 nucleosome, but the acetylation at nucleosome flanking the transcription start site (TSS), did not change substantially (Fig. 5A). Similar changes in H3K27ac patterns were observed across all model genes (Fig. S10), despite wide differences in bursting and expression changes (Fig. 4C-4E). This lack of correlation points to the possibility that bursting rates might not be tightly correlated with promoter H3K27ac levels in agreement with recent reports^39, 53, 60^. In support of uncoupling of promoter H3K27ac and gene bursting behavior, we also did not see any correlation between the native TS frequency and the integrated H3K27ac ChIP signal in DMSO controls at the promoters of all analyzed genes (TSS±1 kb, Fig. 5B). Similar lack of correlation was also seen at the TSS upstream region (either 1kb or 0.5kb). Finally, changes in TS frequency with panobinostat were also uncorrelated with promoter H3K27ac changes (Fig. 5C). Taken together, our results indicate that transcriptional bursting rates are stoichiometrically uncoupled from promoter H3K27ac levels.

**Figure 5:**
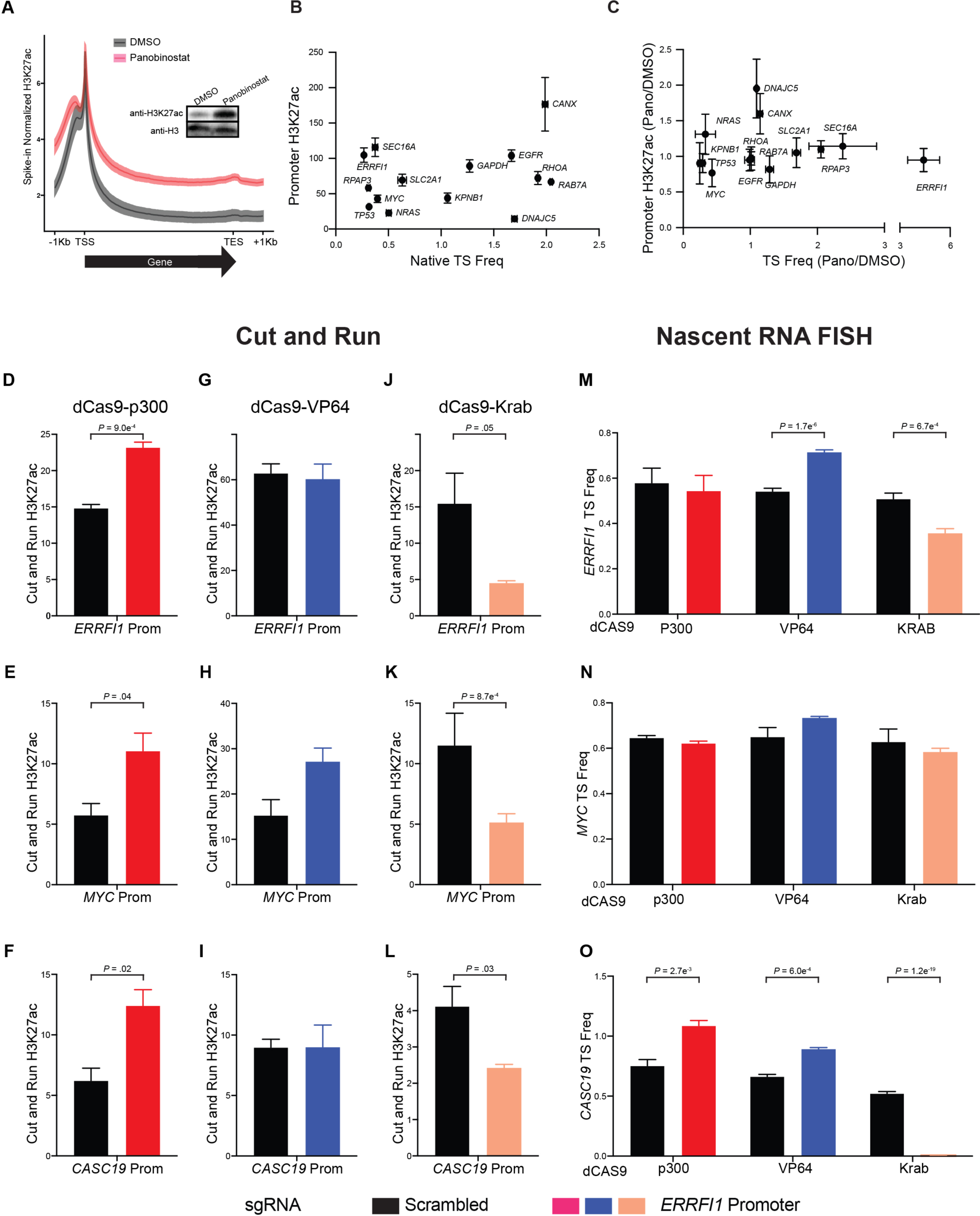
Changes in TS frequencies are stoichiometrically uncoupled from chromatin acetylation changes. **A**. Meta-gene plot of H3K27ac levels at the gene-body ± 1kb region in HBEC treated with panobinostat or DMSO (control) for 4 h. H3K27ac levels between samples were normalized using spike-in *Drosophila* chromatin for comparison. The values represent mean ± SEM from at least 2 and 3 biological replicates for control and panobinostat treated cell, respectively. The inset shows the western blot for total H3K27ac and H3 loading control in treated and control cells. **B** and **C**. Plots depicting the lack of correlation between native TS frequency and H3K27ac levels at the promoter region (TSS ± 1kb) in (**B**) DMSO controls and (**C**) the relative changes in panobinostat (Pano) vs DMSO control for model genes used in the screen. Values represent mean ± SEM from at least 2 biological replicates. **D-O**. Spike-in normalized qPCR coupled Cut-and-Run assay for promoter H3K27ac levels (**D-L**) and nscRNA-FISH (**M-O**) for *ERRFI1* (**D**, **G**, **J** and **M**), *MYC* (**E**, **H**, **K** and **N**) and *CASC19* (**F**, **I**, **L** and **O**) upon dCas9 tethering of p300 (**D- F**), VP64(**G-I**) or Krab(**J-L**). Cut-and-Run values represent mean ± SEM of at least three biological replicates. TS frequencies in **B**, **C**, **M**, **N** and **O** were estimated from >300 cells for each experiment. *P* value (t-test) for the significant differences from the controls are indicated.

To more rigorously test the causality, or lack thereof, between promoter H3K27ac and bursting behavior, we tethered the C-terminal catalytic portion of the p300 acetyltransferase to the promoters of *ERRFI1*, *MYC* or *CASC19*, a non-coding RNA near *MYC*^61^, using dCas9- tethering^62, 63^ (Fig. 5 D-F; see Methods). We assayed both transcriptional bursting and promoter H3K27ac levels by nscRNA-FISH and by qPCR-coupled spike-in normalized Cut-and-Run assays^64^, respectively. As expected, tethering of p300 substantially increased promoter H3K27ac of all genes (50-100%), but a significant increase in TS frequency occurred only for *CASC19* (Fig. 5D-F, M-O). This observation agrees with recent reports where the impact of p300 on expression was both gene- and cell-specific^53, 54^. The lack of causality was also evident when we analyzed changes in promoter histone acetylation upon gene-specific changes in bursting. To do so, we tethered the transcriptional activator VP64 or the transcriptional repressive Krab (Krüppel associated box) domain to the gene promoters via dCas9^63^. As expected, VP64 significantly increased TS frequency for *ERRFI1* (40%) and *CASC19* (30%) but did so without significant changes in promoter H3K27ac levels (Fig. 5G, 5I, 5M and 5O). While *MYC* TS frequency was moderately upregulated (8%), it showed a ∼90% increase in promoter H3K27ac (Fig 5H and 5N). On the other hand, Krab tethering decreased TS frequency from 10% for *MYC* to 90% for *CASC19* but promoter H3K27ac decreased more than 50% for all three genes (Fig. 5J-5L, 5M-5O). Overall, we conclude that promoter H3K27ac levels are uncoupled from bursting behavior and as such are unlikely a dominant determinant of transcriptional bursting patterns.

### Acetylation of Integrator complex regulates bursting kinetics

Our data suggests a role of acetylation in regulating several aspects of bursting, but not a clear deterministic function of promoter histone acetylation. An alternative mechanism is via acetylation of non-histone proteins. Indeed, a wide range of cellular proteins including transcriptional regulators are acetylated^65–67^. To explore this possibility, we performed tandem mass spectrometry to identify differentially acetylated nuclear proteins upon deacetylase inhibition. Isolated nuclei from control (DMSO) and 4 h panobinostat treated HBECs were used to generate libraries of tryptic peptides that were barcoded and immunoprecipitated with antibody to acetylated lysine before tandem mass spectrometric detection^68, 69^. A total of 57 and 46 proteins were found to have acetylated lysines in the two independent replicates, out of which 43 and 33 proteins, respectively, showed an increased acetylation with panobinostat (Table S5). Stringent filtering of the shared hits that included selection of hits that were robustly detected in both DMSO and panobinostat treated cells and showed reproducible enrichment upon treatment with panobinostat, yielded a set of eight proteins (see Methods; Fig. 6A). In agreement with previous studies, these hits included histones *H1*, *H3.3*, *H4* and *H2B,* validating our approach^65, 70^. In addition, we identified four non-histone proteins with unrelated functions: *HMGA1*, *INTS4*, *NCL*, and *RPL14*. Given that increased global acetylation results in pervasive bursting changes across cell types, we hypothesize that at least some non-histone targets of acetylation with a known role in transcription will be conserved between cell lines. Thus, to both rigorously test our methodology and to identify conserved hits, we repeated our mass spectrometric assay in HCT116 cells and found all non-histone hits in HBEC were conserved. Overall, 190 proteins contained one or more acetylated lysine of which 130 showed increased acetylation upon panobinostat treatment, including many histones (Fig. S11 and Table S5). The major biological processes represented in the gene ontology analysis of these hits relative to all 2004 proteins detected in the nuclear lysate included regulation of transcription by Pol II, chromatin organization/remodeling and regulation of RNA biosynthesis. Thus, hits from both cell lines suggest that many nuclear proteins, besides histone, are acetylated and amongst these hits several regulators of transcription show increased acetylation with deacetylase inhibition. Out of the common non-histone hits we choose to study the role of acetylation at lysine 26-27 of Integrator complex subunit 4 (*INTS4*) on gene bursting in HBEC due to its direct role in regulating late steps in transcription^71^.

**Figure 6:**
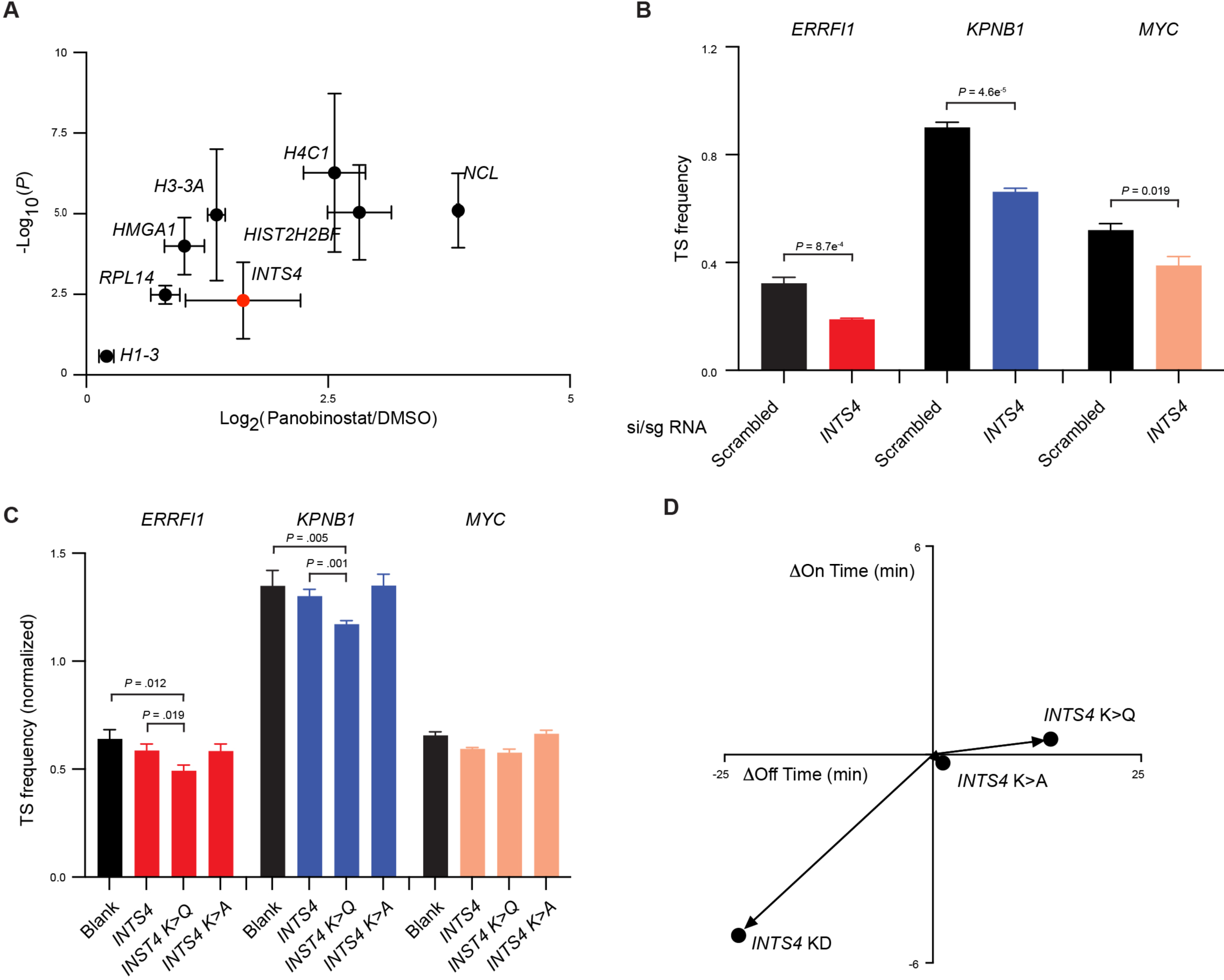
Acetylation at Ints4 K26-27 decreases bursting of target genes. **A**. Enrichment plot for genes based on the increased levels of immune-enriched acetylated lysine containing peptides (Log_2_FC>0.2) derived from nuclei of HBEC treated with panobinostat for 4 h with respect to control treatment (DMSO). The values represent mean ± SEM from two experiments with 5 and 9 biological replicates. **B**. TS frequency for *ERRFI1*, *KPNB1* and *MYC* after 48 h depletion of *INTS4* using both siRNA and sgRNA targeted to *INTS4* gene body and promoter, respectively, or scrambled controls in wild-type cells constitutively expressing dCas9-*KRAB*. **C-D**. Assaying the effect of overexpressing wild-type and mutant *INTS4*. TS frequencies for *ERRFI1*, *KPNB1* and *MYC* in monoclonal wild-type HBEC lines with no ectopic *INTS4* (Blank) or ectopic expression of either wild-type *INTS4*, acetylation mimic (*INTS4 K>Q*) and acetylation null (*INTS4 K>A*) mutant, normalized by allele frequency relative to HBEC with no ectopic *INTS4*. Both K26 and K27 were altered in the mutants. The TS frequency values in **B** and **C** represents mean ± SEM of at least four biological replicates with >300 cells for each experiment and *P* value (t-test) for the significant differences from the controls are indicated. **D**. Mean changes in the Off- and On- times of *ERRFI1* bursting in *INTS4* depleted cells or monoclonal cells over expressing the *INTS4* mutant (acetylation mimic and null), with respect to the scrambled control or cells overexpressing wild- type *INTS4*, respectively. The mean values were derived from >40 cells from one or more acquisitions.

Ints4 is a scaffolding subunit in the endonuclease subcomplex (Ints4-Ints9-Ints11) of the metazoan-specific and essential RNA processing complex referred as Integrator^72^. Initially identified as 3’ cleavage and processing enzyme for snRNAs, Integrator has recently been shown to cleave 5’ ends of mRNAs^73^. Depletion of the endonuclease subunit (Ints11) results in an increased pause release but inefficient elongation^71, 74^, leading to higher expression of small genes (<5kb) and repression of larger genes^71, 74^. To test for a potential role in gene bursting, we depleted *INTS4* by combining siRNA and CRISPRi and assessed bursting changes by nscRNA-FISH (Fig. 6B and S12A; see Methods). *INTS4* depletion significantly decreased the TS frequency for all genes tested (*ERRFI1*, *KPNB1*, *MYC*; Fig. 6B). Live-cell assay for *ERRFI1* bursting confirmed this effect and indicated that the decrease in bursting was primarily driven by a decrease in On-time (45%), despite a concomitant decrease in Off-time (35%; Fig. 6D). These results confirm involvement of Integrator as a regulator of gene bursting.

To directly test whether the observed effect of protein acetylation on bursting behavior occurred via acetylation of *INTS4*, we generated single cell clones of HBECs that stably expressed comparable levels of wild-type or *INTS4* mutants that either mimic (K26-27Q) or are null for acetylation (K26-27A) (Fig. S12B). Given that Integrator function is essential for cells^75, 76^, ectopic *INTS4* variants were expressed in wild-type background. After normalization for allele frequency to eliminate potential cell cycle effects of *INT4* expression (Figure S13A, B; Table S6), we find that neither the wild-type nor the acetylation null-mutant *INTS4* affected the TS frequency of *ERRFI1*, *KPNB1* or *MYC*. However, in line with the observed reduction of bursting frequency of many genes upon deacetylase inhibition in our initial screen, the acetylation mimic *INTS4* decreased the TS frequency for *ERRFI1* and *KPNB1* genes by 10%-20% and to a lesser extent for *MYC* (Fig. 6C). We validated these changes in live-cell bursting assays. Consistent with nscRNA-FISH (Fig. 6C), the acetylation null mutant exhibited little change, while the acetylation mimic showed a decrease in bursting relative to the wild-type *INTS4* expressing clones (Fig. 6D). The latter resulted primarily from a ∼30% increase in OFF-time (Fig. 6D). We conclude that the Integrator complex promotes gene bursting, and this regulatory event is mediated by the acetylation at K26-27. These results demonstrate a causal role of non-histone acetylation in regulating transcriptional bursting on native genes.

## Discussion

We have developed a high-throughput imaging pipeline for analyzing gene bursting behavior of endogenous genes at the single allele-level. We applied this method for screening a small molecule library of epigenetic modulators to uncover regulatory principles and molecular mechanisms of gene bursting. We find a prominent role of protein acetylation in altering the bursting behavior of multiple endogenous genes. Mechanistically, these effects were not due to promoter-associated H3K27 acetylation. Instead, we provide evidence for an acetylation-dependent regulatory role of the Integrator complex in control of gene bursting, pointing to downstream regulation of gene bursting.

Gene bursting is an evolutionary conserved feature of transcription^6^. Widely distinct patterns of bursting have been observed for individual genes^11^, but the molecular determinants of bursting patterns in general and of gene-specific On- and Off-times are not well understood. This is in part because of experimental limitations due to the use of either engineered reporter system or single-cell RNA-seq approaches that make systematic analysis of endogenous genes challenging^36^. Consequently, analysis of gene bursting often does not fully reflect the complexity in bursting responses. For example, tethering of p300 at the *FOS* promoter in mouse cortical neurons increases On-time, while its tethering at the *BMAL1* promoter in HEK293T cells decreases Off-time^20, 21^. Similarly, global changes in acetylation have been reported to change either Off- or On-time or both for different model genes^11, 18, 19, 40^. To overcome these limitations, we have developed an RNA-FISH-based approach to measure gene bursting patterns through sensitive and robust detection of nascent RNA at individual TS of endogenous genes. We adapted this FISH-based bursting assay to a high-throughput format which allowed screening of a library of small molecule modulators of chromatin modifying enzymes against multiple endogenous genes in several cell lines (Fig. 1A). We focused on fast-acting small molecule modulators, rather than long-term alteration of epigenetic states by RNAi or CRISPR, to assess acute effects on bursting and to limit secondary effects. Using this strategy, we tested the bursting behavior of 14 genes in three cell lines in a total of 5,880 experimental conditions (Fig. 2A, 3A and S7A). Because of its focus on endogenous genes and its high-throughput format, our approach offers the opportunity to uncover general principles of gene bursting changes upon acute changes in chromatin states.

Using TS frequency and intensity as read outs, the combined screening data highlighted two key principles underlying gene bursting (Fig. 2A and 3A). First, the native bursting rate of a gene strongly affects both the magnitude and nature of bursting changes in response to perturbation. Specifically, we show that the degree of change in TS frequency is inversely related to the native TS frequency of genes (Figure 2B). Notably, none of our perturbations increased the burst frequency for the most highly bursting genes, suggesting there is a refractory period or an upper limit of transcriptional bursting that is not readily surpassed. Furthermore, the nature of the bursting change, i.e. alterations in frequency and intensity of TS, is also sensitive to the bursting rate as illustrated by the fact that the frequency and intensity changes are concomitant for genes with intermediate bursting rates (TS frequency = 0.5-1.5), but largely independent for low and high bursters (TS frequency < 0.5 and >1.5; Fig S6C). Second, we find that the impact of epigenetic changes on bursting is not universal, but rather depends on both the gene- and cell-line context. Thus, two genes in the same cell or the same gene in two cell lines can respond in diametrically opposite fashion to the same perturbation. For example, most deacetylase inhibitors upregulated *ERRFI1* and downregulated *MYC* bursting in HBEC and MCF7, but the same set of inhibitors decreased bursting of both genes in HCT116 (Fig. 2A). These results explain some of the previously reported variability of effects in response to individual perturbations on different genes and in different cell lines^11, 18–21^. They also highlight the need for systematic, rather than anecdotal, analysis of the effect of chromatin states on gene bursting.

We chose to study the effect of acetylation on gene bursting in more detail because amongst all classes of epigenetic modulators tested those that targeted deacetylases, bromodomain proteins and acetyltransferases led to the biggest changes in TS frequency, intensity and heterogeneity (Figure 2C, 3C and S7C). Live-cell bursting assays on multiple model genes that cover a broad spectrum of bursting rates and expression levels revealed an almost uniform decrease in Off-times in response to protein acetylation changes (Fig. 4C). While this finding is in line with previously reported increases in burst frequency upon increased acetylation^11, 19, 20, 40^, we have now shown that both the directionality and magnitude of On-time changes are highly gene-specific and that they determine the overall changes in bursting and expression (Fig. 4D and 4E). For example, of five genes tested only *ERRFI1* with a 57% increase in On-time showed both an increase in TS frequency and expression, while a decrease in On-time for other genes by 15-47% led to downregulation to different extents despite a >25% decrease in Off-times (Fig. 4C). These results provide a kinetic basis for the diverse responses to global increase in acetylation reported previously^60, 70^.

We also tested the mechanistic role of histone acetylation on gene bursting in more detail by comparing bursting behavior of multiple genes with promoter-proximal H3K27ac, the acetylation mark most commonly associated with active transcription^52^. A clear correlation was absent in the native states or upon perturbations by small molecules and upon tethering-induced gene-specific changes in promoter H3K27ac (Fig. 5). While our results suggest that promoter-associated H3K27ac is not a major driver of bursting, they do not rule out roles for H3K27ac or other histone acetylation marks in bursting regulation. For example, H3K27ac and other acetylation marks appear to play a role in maintaining the bursting state, which is consistent with the role of H3K27ac as an anti-repressive mark^77^. Alternatively, histone acetylation may promote bursting in the context of only some chromatin states, which is consistent with our observation that only some, but not all, genes exhibit increased bursting upon promoter-specific acetylation (Fig. 5D-5O). Furthermore, other acetylation marks such as H4K16ac, which affects nucleosome stability, may promote bursting^78, 79^. Although, several lines of evidence indicate that acetylation-related gene expression changes are not a direct result of changes in nucleosome occupancy^80, 81^. Taken together, these results point to a role of non-histone protein acetylation in gene bursting.

We indeed identified subunit 4 of the Integrator complex (*INTS4*) as a prominent non-histone acetylation target in multiple cell lines and demonstrate its role in bursting regulation (Fig. 6A and S11). *INTS4* codes for a scaffolding protein in the endonuclease subcomplex and contains key acetylation sites, K26 and K27, in its flexible N-terminal tail^82^. Consistent with a global role in transcription regulation we find that modulation of *INTS4* levels or mutation of its acetylation sites alters bursting (Fig. 6B-6D). While over-expression of the acetylation-mimic *INTS4* decreased bursting significantly, little change in bursting was observed with similar overexpression of either wild-type or acetylation mutant (Fig. 6C and 6D), suggesting a repressive role for acetylation of Ints4 in determining bursting patterns. One possible mechanism could be that acetylation of Ints4 alters endonuclease activity by altering the interactions within the Ints4-Ints9-Ints11 subcomplex, especially given that acetylation of non-histone proteins may alter protein-protein interactions^66, 83^. Alternatively, the relative levels of the endonuclease subcomplex containing either Ints4 or Brat1, a recently identified scaffolding subunit of endonuclease subcomplex^84, 85^, may change with Ints4 acetylation. While we have identified a causal role of Ints4 acetylation on bursting, Ints4 acetylation does not explain all the bursting changes we see with deacetylase inhibition. Interestingly, we find in our mass-spec analysis several other regulators of transcription that are significantly acetylated upon deacetylase inhibition (Figure S11, Table S5). Future investigation into the role of these acetylation targets on bursting will shed more light on the relative contributions of histone modifications and transcription factors in gene bursting.

Taken together, our results provide novel insights into the molecular determinants of gene bursting. We identify a strong relationship between the sensitivity of a gene’s bursting pattern and its basal bursting properties. Furthermore, our finding of an acetylation-mediated regulatory role of the Integrator complex points to far-downstream regulation of gene bursting via control of transcription-pause release.

### Limitations of the study

Our study has some technical limitations. While nscRNA-FISH detection is robust, we cannot exclude the possibility that alleles with very low expression are missed in our analysis. Similarly, accurate measurements of FISH signal intensities are notoriously challenging. These concerns are partially mitigated by the fact that all our analyses are comparative in nature and inaccuracies in TS detection or intensity would most likely only affect the extent of any observed changes. A further limitation of our approach are potential off-target effects of the small molecule inhibitors used, although we see consistent behavior amongst multiple inhibitors of the same class for the same target. With regards to mass spectrometric studies, it is possible we missed low abundance targets and the number of hits we obtained may be sensitive to day-to-day variation in sample preparation. Finally, we noticed in our studies that the effects of overexpression of Ints4 were somewhat dependent on the expression level and upon high expression slowed down the growth rate. We consequently limited our studies to use of cell lines with similar and relatively low expression levels (Fig. S12B).

## Acknowledgments

We thank the members of the Misteli lab and Larson lab for their feedback on the manuscript. We thank the members of the core facility at CCR for high-throughput imaging, deep-sequencing, flowcytometry and high-performance computing group. This research was funded by grant #1ZIABC010309-24 from the Intramural Research Program of the NIH.

## Author contributions

V.S., T.M. and D.R.L conceptualized the project, designed the experiments and interpreted the results. T.M. obtained funding. V.S. executed experiments, R.H. performed mass spectrometry experiment under T.A.s guidance. V.S., T.M. drafted the manuscript. V.S., T.M. and D.R.L participated in reviewing and editing the manuscript.

## Supplemental information

**Figure S1:**
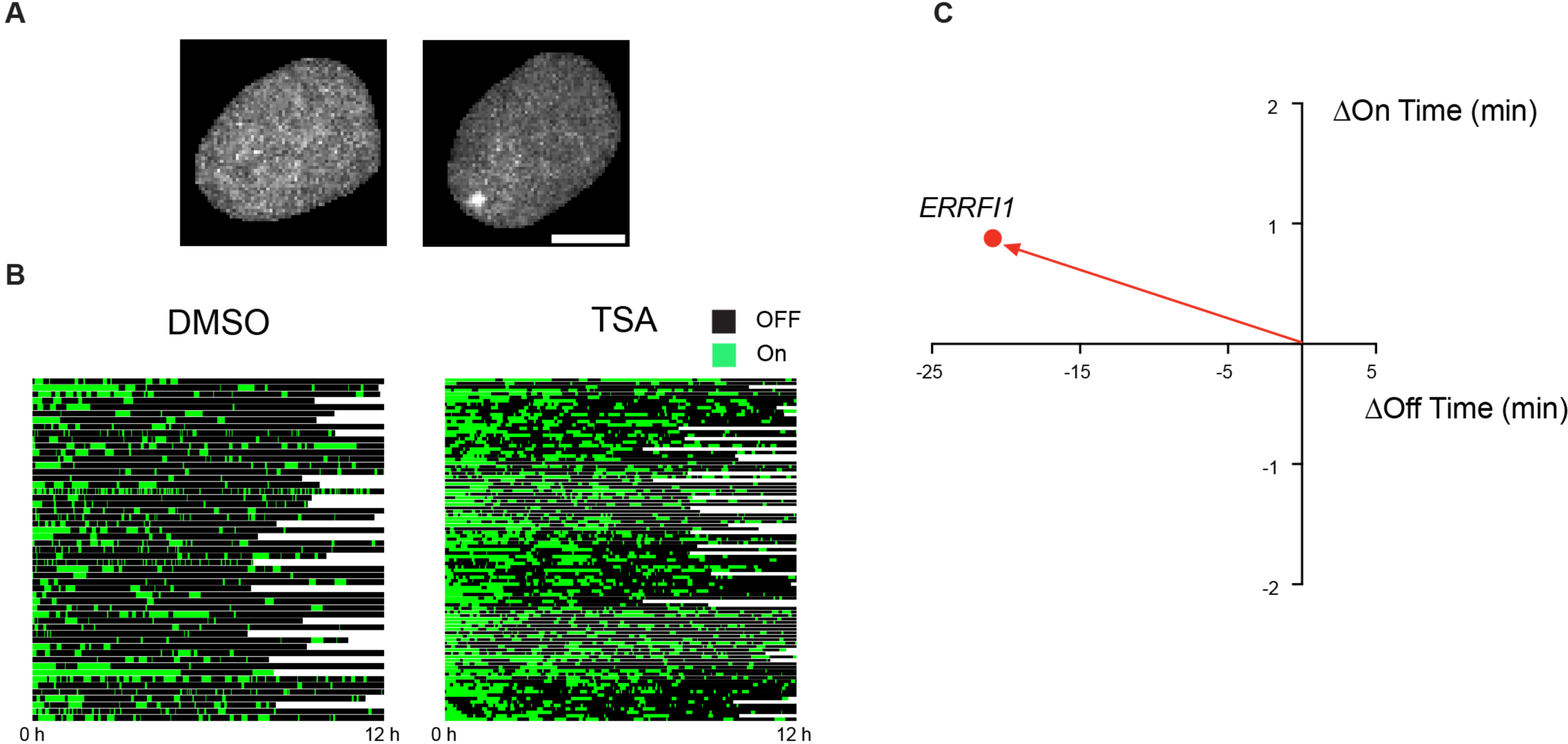
Live-cell bursting of *ERRFI1* with trichostatin. **A.** Representative images of HBEC nuclei with monoallelic intron insertion of MS2 operator array in *ERRFI1* and over expressing MS2-GFP during Off- (left) and On- (right) states. The scale bar represents 5μm. **B** and **C**. Bursting kinetic changes for *ERRFI1* in trichostatin A (TSA). Cells were treated with either 1μM TSA or DMSO for 4 h and *ERRFI1* bursting imaged for next 12 h at 100 sec intervals. Both cells and MS2-GFP spot were segmented and tracked (See methods). **B**. Kymogram for *ERRFI1* bursting. Each row represents a single trace of MS2-GFP spot intensities over time for an allele. **C**. Changes in the mean Off- and On-times of *ERRFI1* bursting. The mean values were derived from >40 cells from one or more acquisitions.

**Figure S2:**
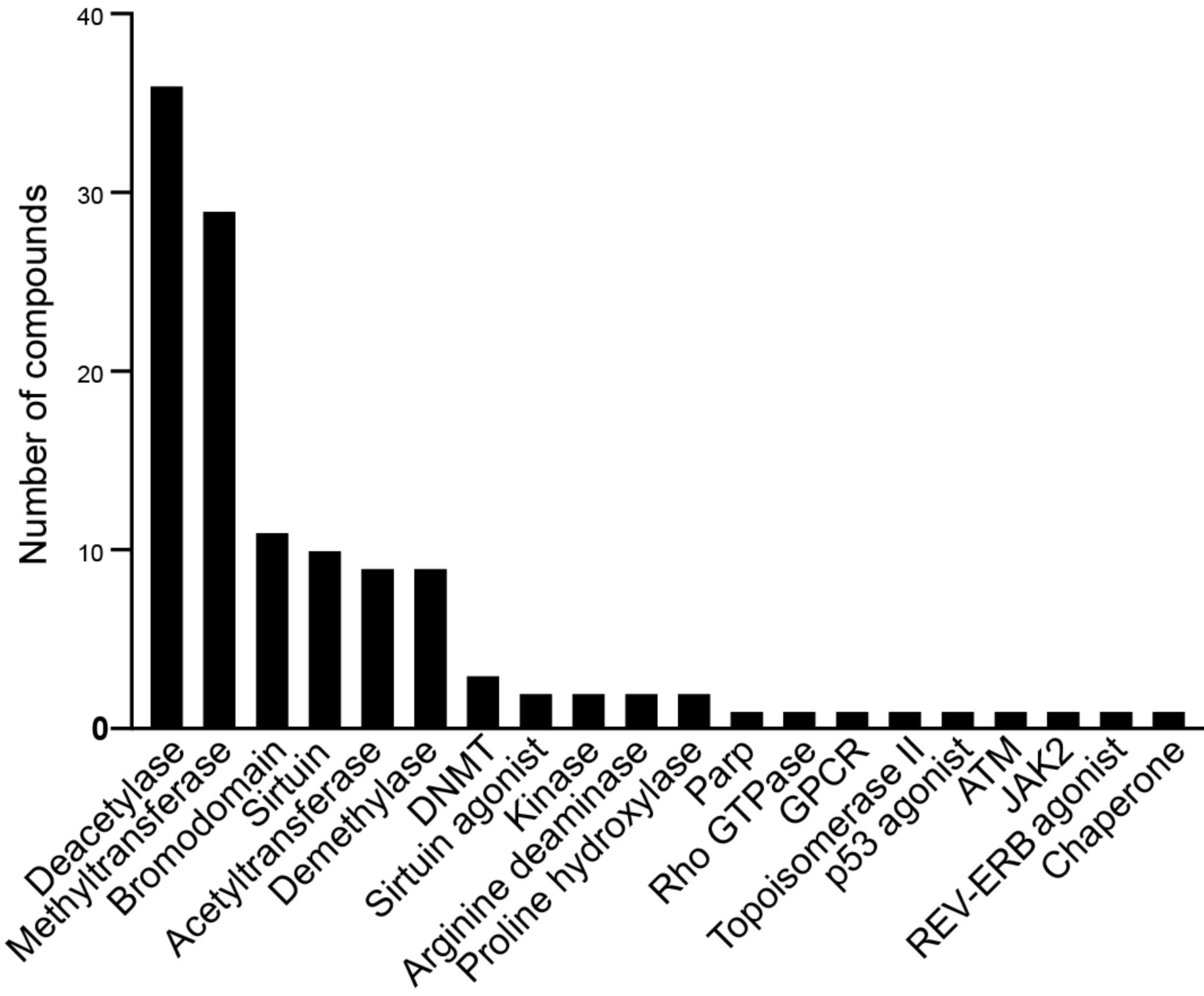
Chromatin modifying enzyme classes targeted in the screen. Frequency distribution of the small molecule modulators for the 20 classes of chromatin modifying enzymes targeted in the screen.

**Figure S3:**
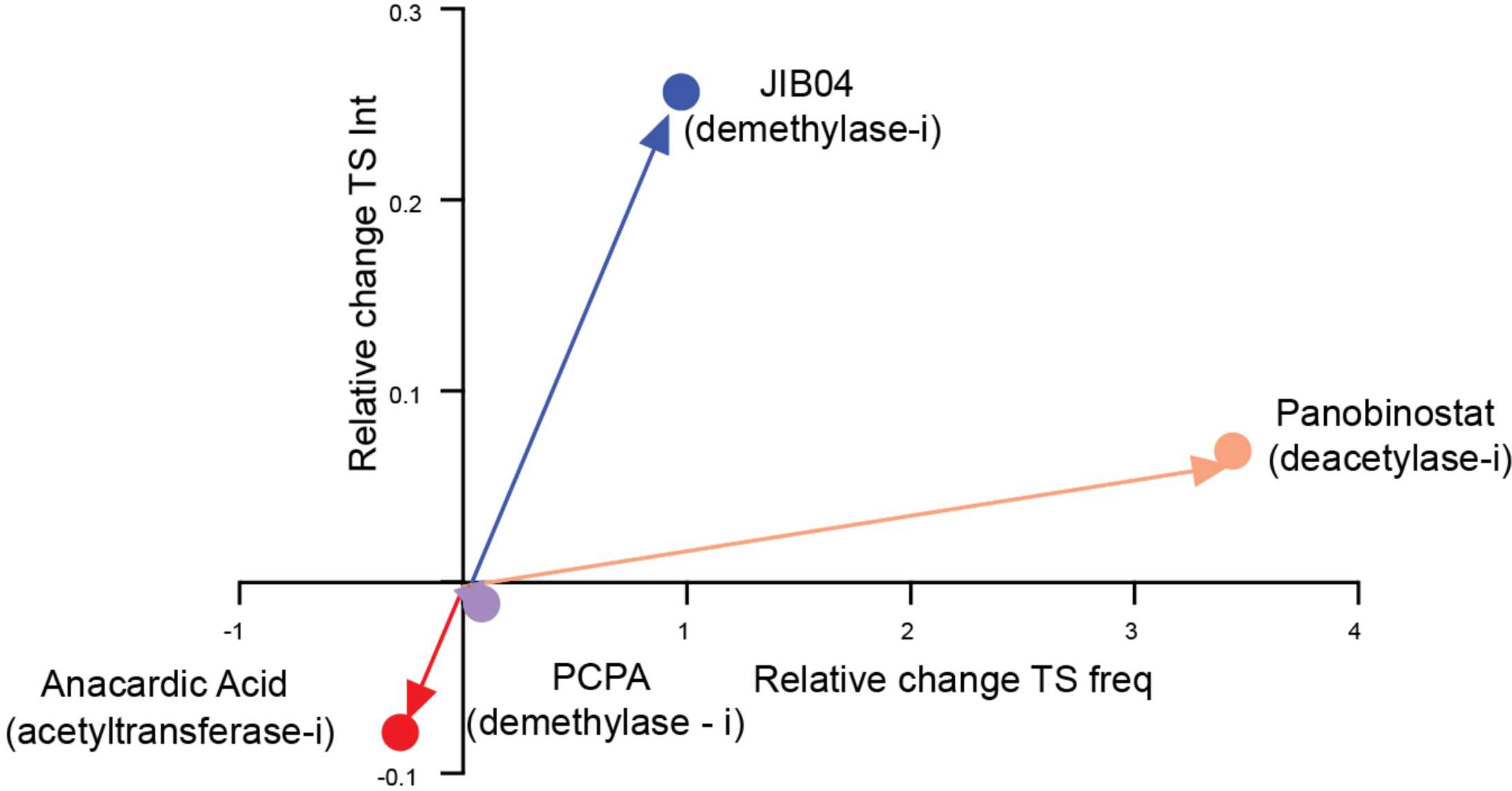
Changes in *ERRFI1* TS frequency and intensity reflect changes in live-cell bursting kinetics. Relative changes in *ERRFI1* TS frequency (Freq) and intensity (Int), (Freq_Treated_- Freq_Untreated_)/Freq_Untreated_ and (Int_Treated_-Int_Untreated_)/Int_Untreated_, respectively, after 4 h of treatment with selected inhibitors used for live-cell bursting assay in Fig. 1E. The TS frequency and intensity values represent mean ± SEM of at least two biological replicates with >300 cells for each experiment.

**Figure S4.**
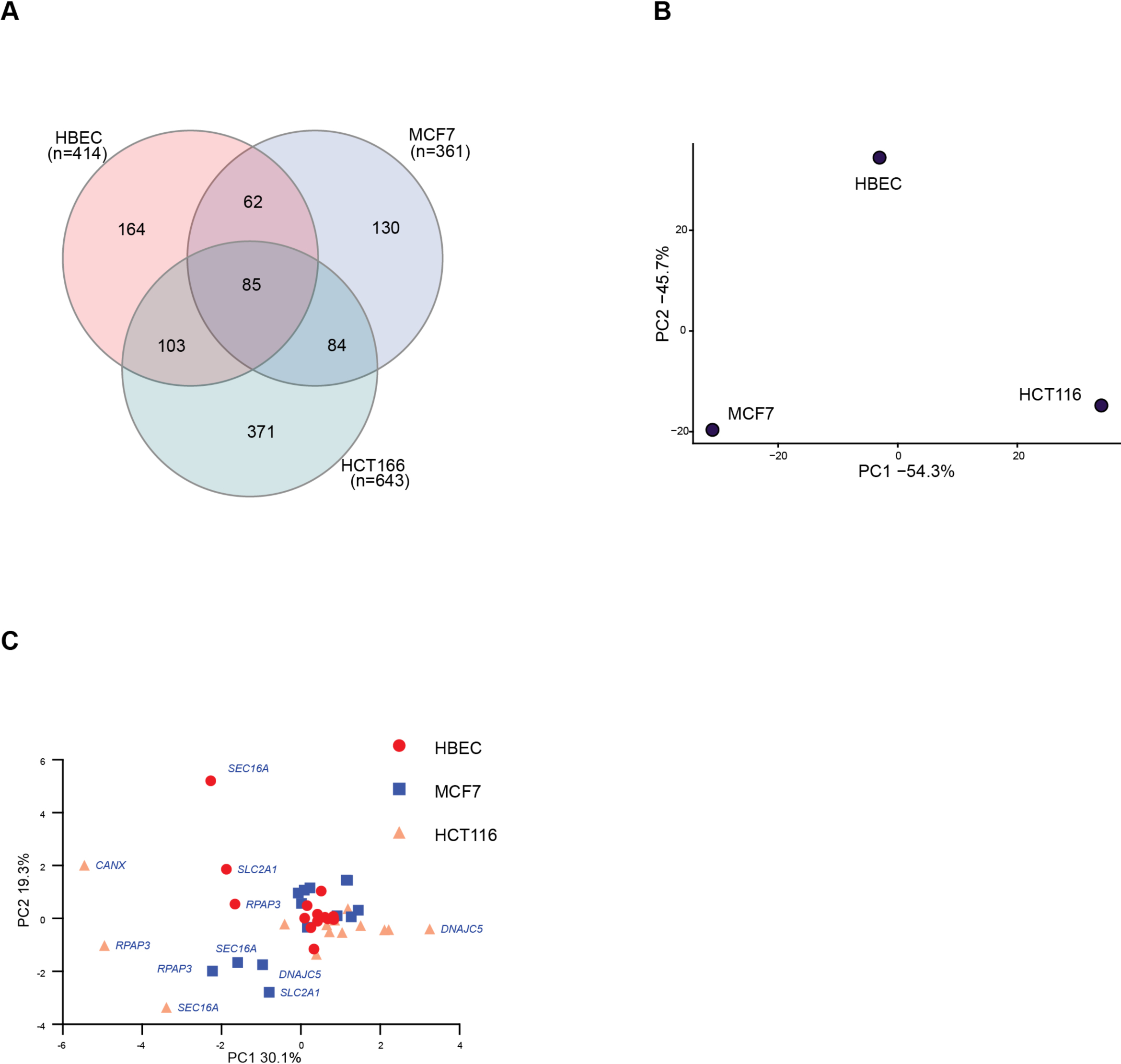
TS frequency changes are weakly conserved across cell lines. **A**. Shared numbers of significantly changed TS frequencies (p<0.05) by small molecule modulators of chromatin modifying enzymes across all model genes between three cell lines: HBEC, MCF7 and HCT116. **B** and **C**. Responses to small molecule modulators of chromatin modifying enzymes are both gene and cell line specific. Principal component analysis of changes in TS frequencies comparing variations between cell lines (**B**) and individual genes (**C**). Genes with higher variations in TS frequency are labelled in blue in **C**. All TS frequency measurements were obtained from at least two biological replicates with >300 cells for each experiment.

**Figure S5:**
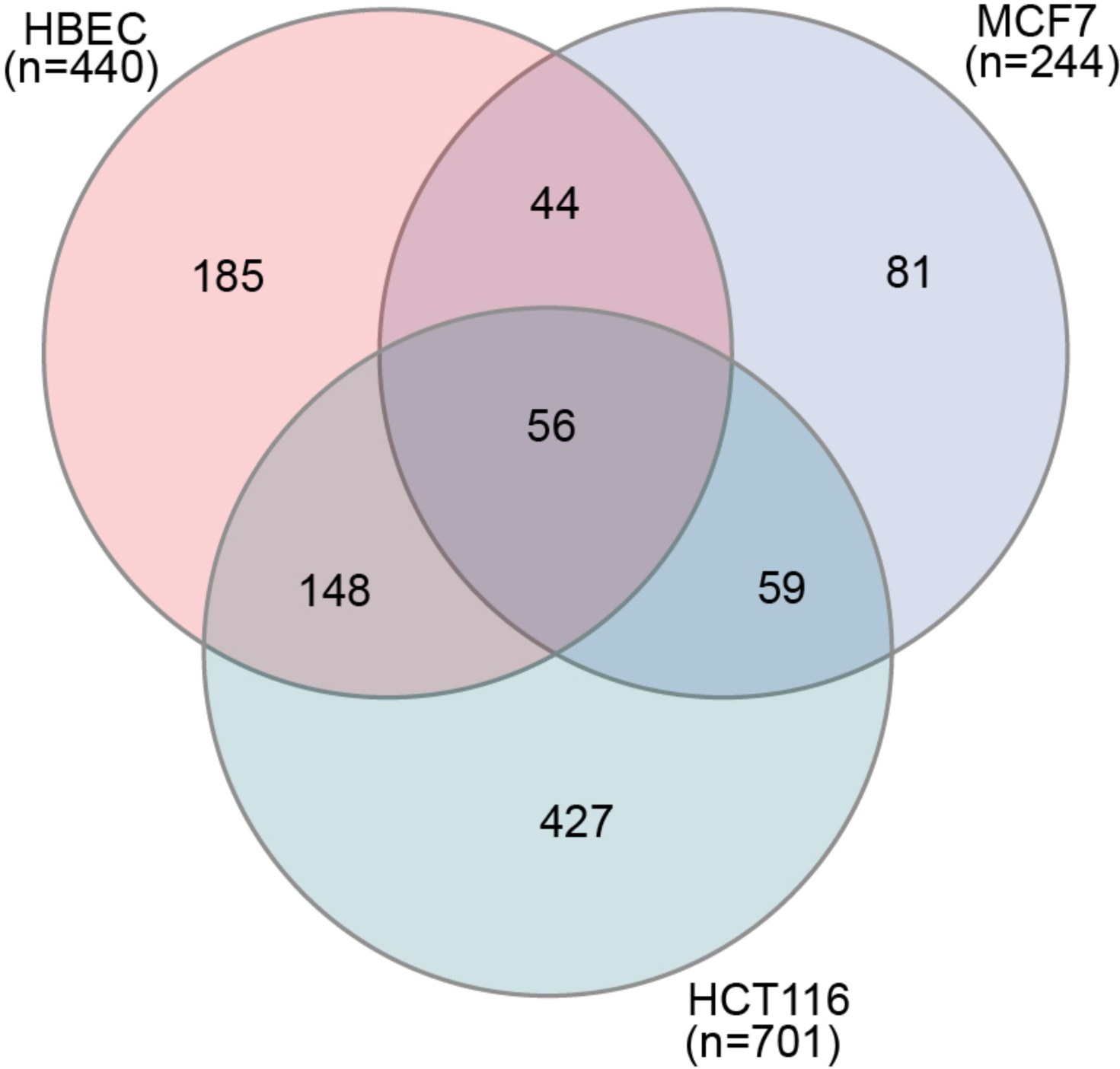
TS intensity changes are weakly conserved across cell lines. Shared numbers of significantly changed TS intensities (*P*<0.05) by small molecule modulators of chromatin modifying enzymes across all model genes between three cell lines HBEC, MCF7 and HCT116. The TS intensity for the hits were obtained from two biological replicates with >300 cells for each experiment.

**Figure S6:**
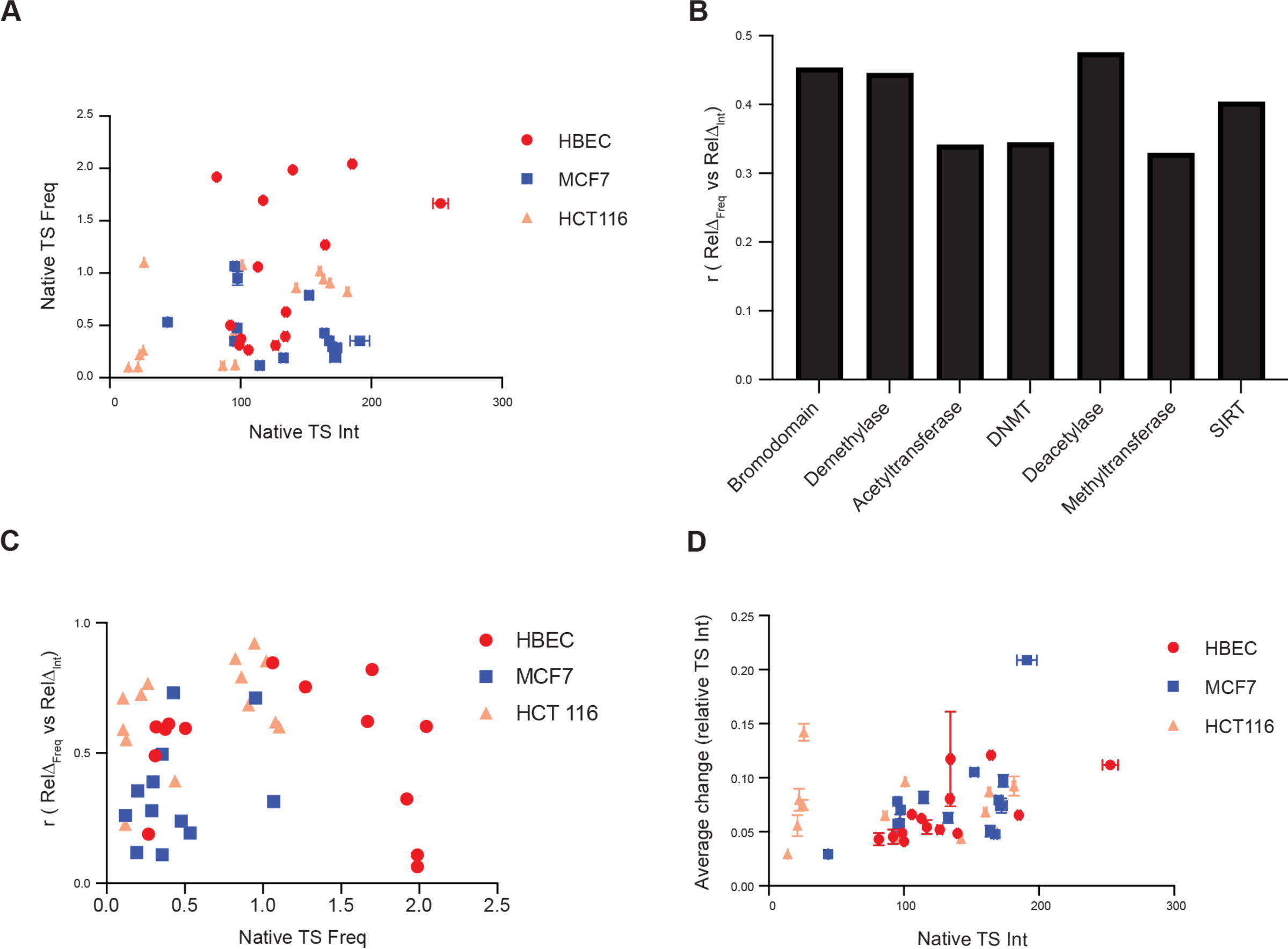
Correlation of TS intensity with other bursting metric from the screen. **A**. Relationship between the native values of TS intensity (TS Int) and TS frequency (TS Freq) for the model genes. The values represent mean ± SEM of at least ten biological replicates of >300 cells. **B** and **C**. Factors affecting the correlation between relative changes in TS frequency (Freq_Treated_- Freq_Untreated_)/Freq_Untreated_) and TS intensity (Int_Treated_-Int_Untreated_)/Int_Untreated_). Spearman rank coefficient (r) of correlation plotted for the major inhibitor classes (**B**) or for model genes with respect to their native TS frequencies (**C**), across three cell lines. The TS frequency and intensity were obtained from at least two biological replicates of >300 cells. **D**. Relationship between the native values of TS Int and the average change in relative TS intensity (Relative TS Int), defined as the average magnitude of TS intensity change relative to the native TS intensity for a gene across the screen, avg(|Int_Treated_-Int_Untreated_|)/Int_Untreated_. Values represent mean ± SEM of at least ten and two biological replicates of native TS intensity and average magnitude change, respectively.

**Figure S7:**
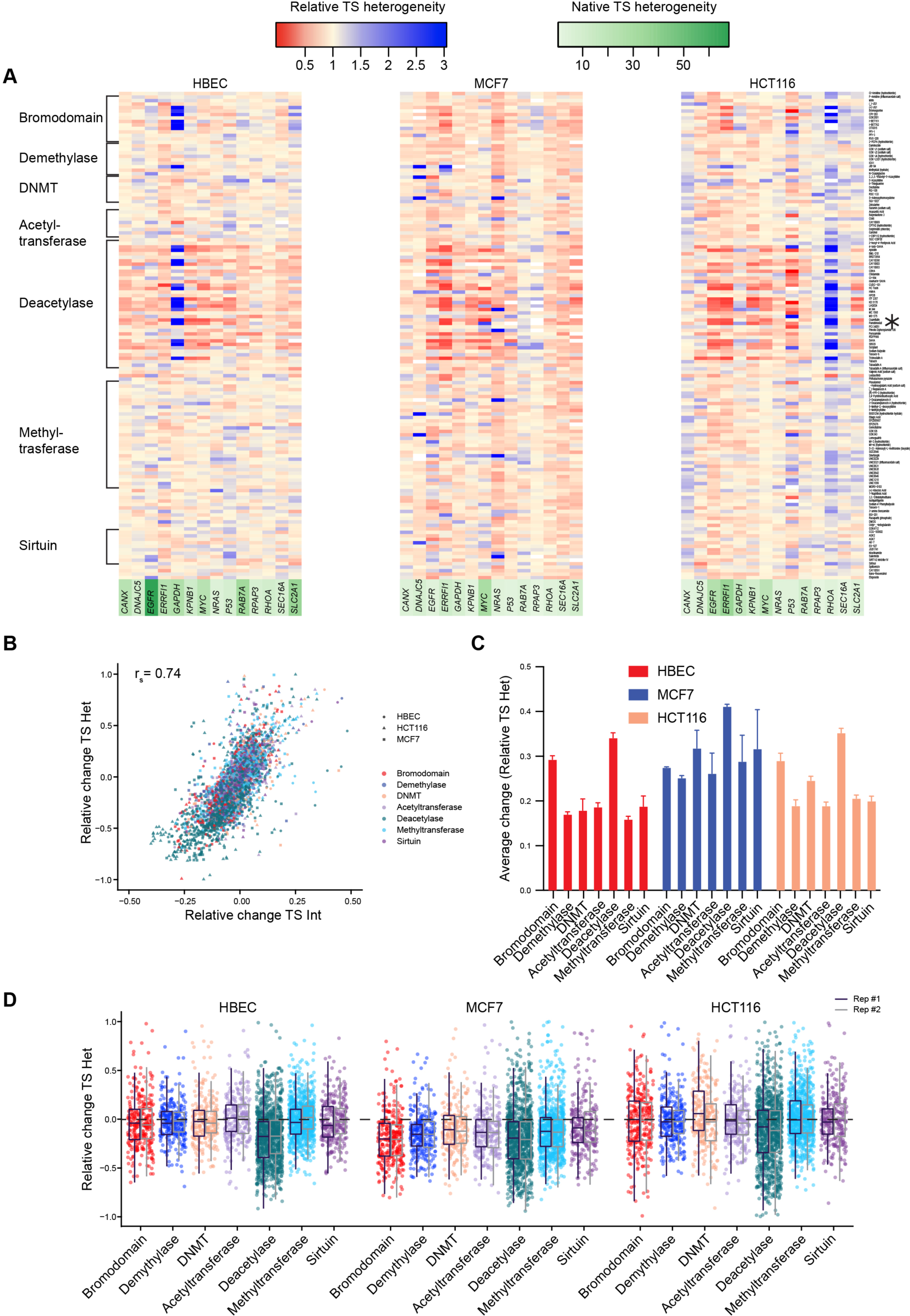
The changes in heterogeneity of TS intensity are most sensitive to acetylation and correlated with changes in burst size. **A**. Heat maps for the relative heterogeneity in TS intensity (Relative TS Het) calculated as the variance over mean of TS intensity (σ^2^/µ) in presences of small molecule modulator targeting different classes of chromatin modifying enzymes relative to the DMSO control for model genes in three cell types: HBEC, MCF7 and HCT116. Treatments with insufficient TS intensities are colored white. The rows are arranged by the modulator class with major classes indicated on the left, panobinostat treatment is asterisked on the right and genes names are color coded based on their heterogeneity in controls. Data represents mean of two biological replicates. **B**. Relationship between relative changes in TS intensities (Int_Treated_- Int_Untreated_)/Int_Untreated_ and the heterogeneity in TS intensities (Het_Treated_-Het_Untreated_)/Het_Untreated_. Spearman rank coefficient (r_s_) indicated. The values represent the mean of two biological replicates between ranges of −1.0 to 1.0 and −0.5 to 0.5 for heterogeneity and intensity changes, respectively. **C** and **D**. Quantitating the impact of the major classes of chromatin modifying enzymes on TS heterogeneity changes. **C**. Comparison of the average change in relative TS heterogeneity, defined as the magnitude of TS heterogeneity changes relative to the native TS heterogeneity averaged across all modulators in the respective classes, avg(|Het_Treated_- Het_Untreated_|)/Het_Untreated_, in three cell lines. Values represent mean ± SEM of two biological replicates. **D**. Relative changes in TS heterogeneity (Het_Treated_-Het_Untreated_)/Het_Untreated_) for each modulator of the respective classes from three cell lines. Values from the two biological replicates are show adjacent. All TS heterogeneity measurements were derived from imaging of >300 cells per condition and data shown for relative change between 1 and −1.

**Figure S8:**
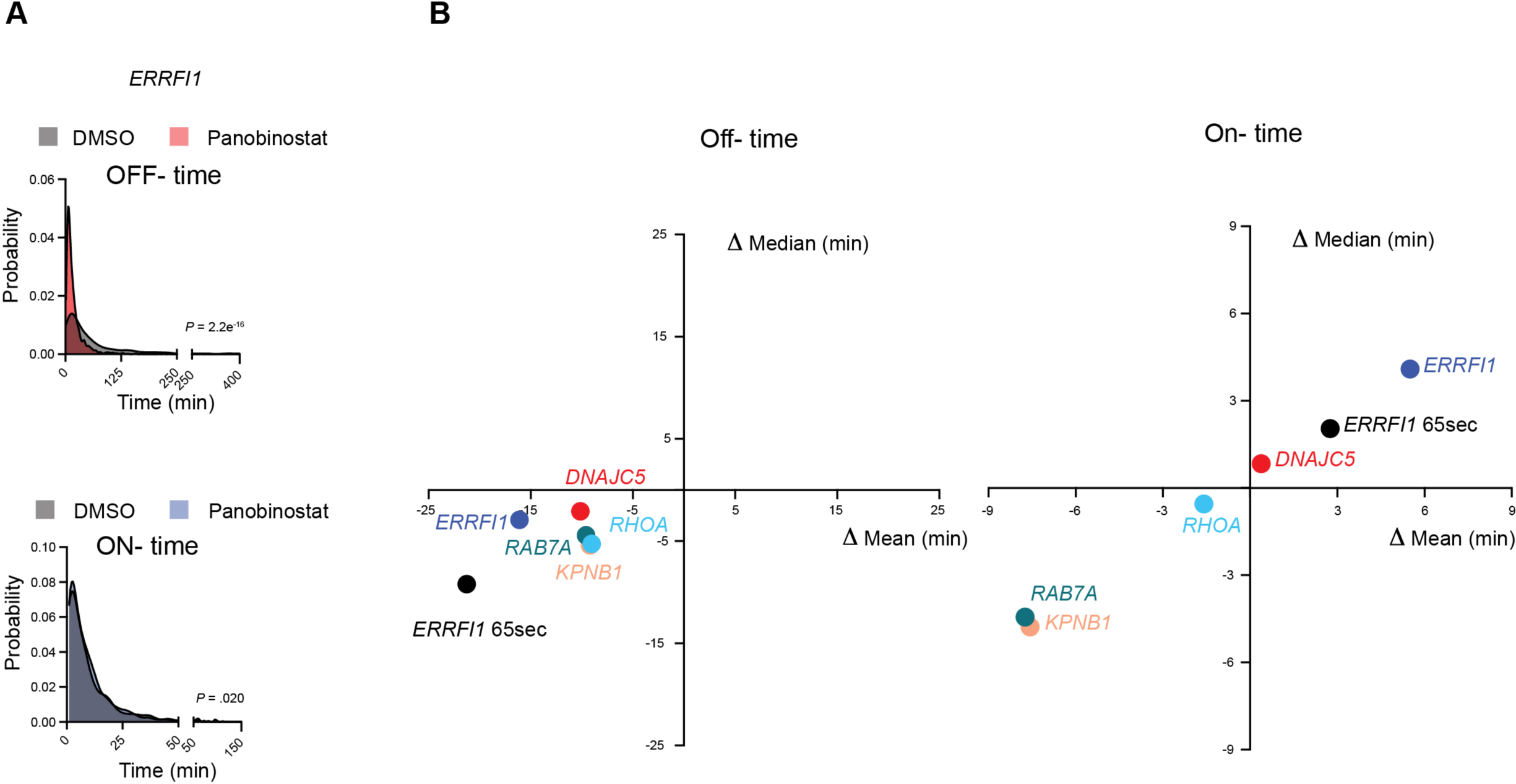
Changes is bursting patterns are consistent at lower acquisition interval. Off-time changes with deacetylase inhibitor (Panobinostat) results from non-uniform changes in the Off- time distribution. **A**. Normalized histograms for the Off- (left) and On-(right) time distribution of *ERRFI1* bursting assayed at 65 s interval in panobinostat treated cells and DMSO control. *P* value: Kolmogorov–Smirnov test. **B**. Comparison of the mean vs median differences in Off- (left) and On- (right) times for model genes upon treatment with panobinostat, relative to DMSO control. Both the distribution in **A** and the mean values in **B** were derived from imaging of >40 cells from one or more acquisitions.

**Figure S9:**
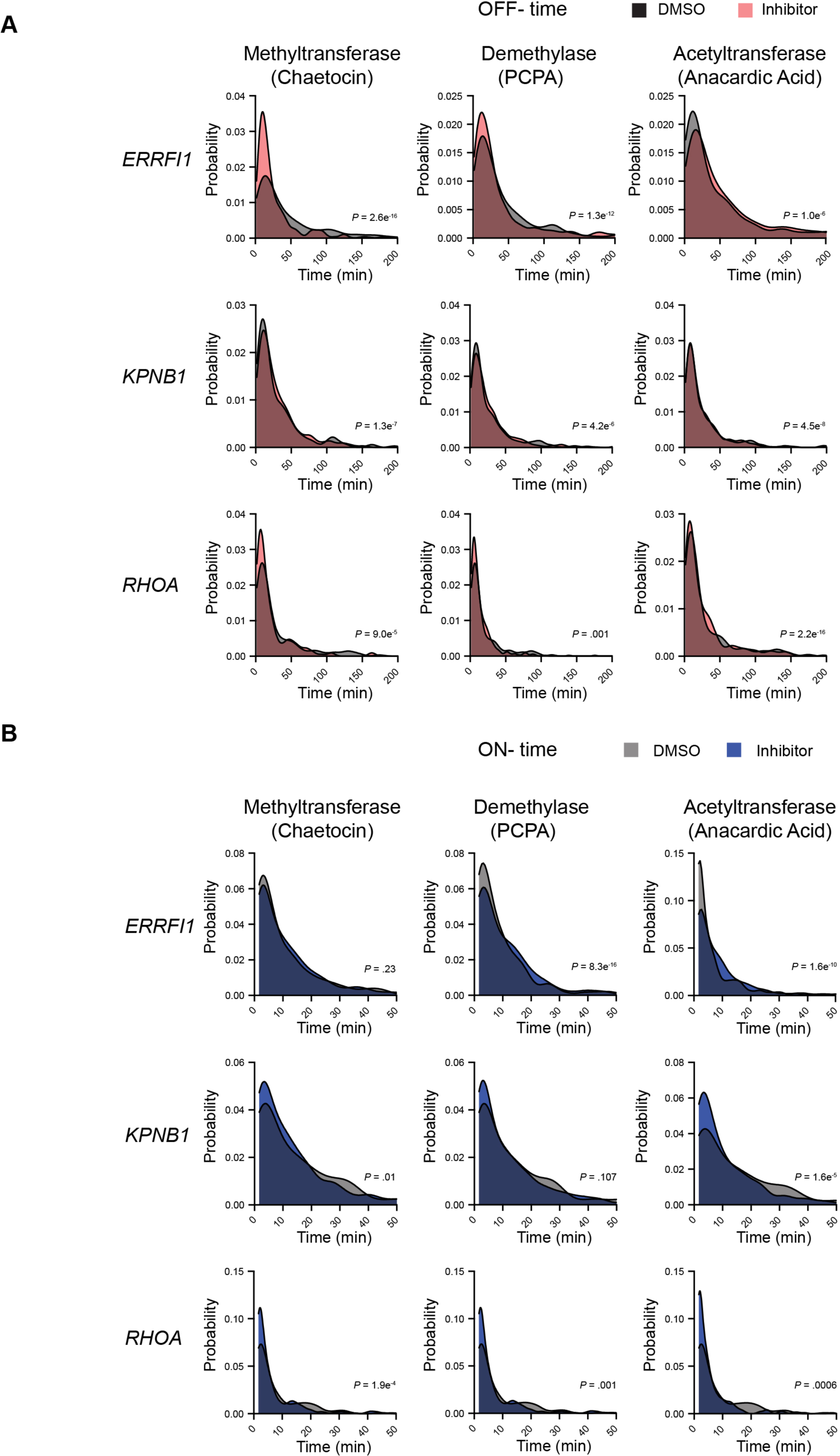
Changes is bursting patterns are similar across inhibitor classes. Normalized histograms for the Off- (top) and On-(bottom) time distribution for bursting of *ERRFI1*, *KPNB1* and *RHOA* in presence of inhibitor targeting methyltransferase (chaetocin), demethylase (PCPA) and acetyltransferase (anacardic acid) or DMSO control. The distributions were derived from imaging of >40 cells from one or more acquisitions. *P* value: Kolmogorov–Smirnov test.

**Figure S10:**
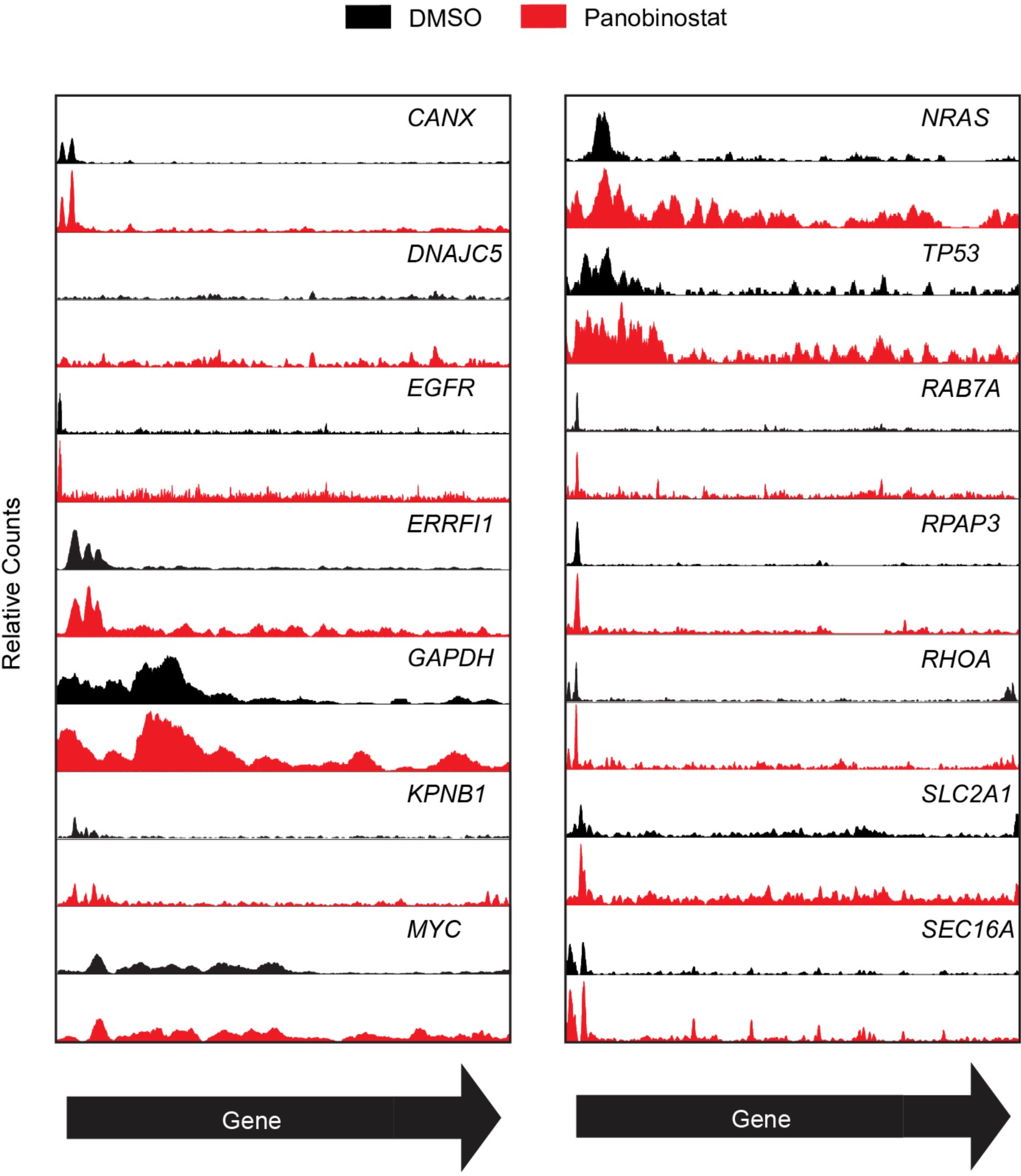
Genes exhibit similar H3K27ac changes with deacetylase inhibitor. Spike-in normalized H3K27ac ChIP-seq tracks in panobinostat treated cells and DMSO control for the gene body ± 1kb region of model genes.

**Figure S11:**
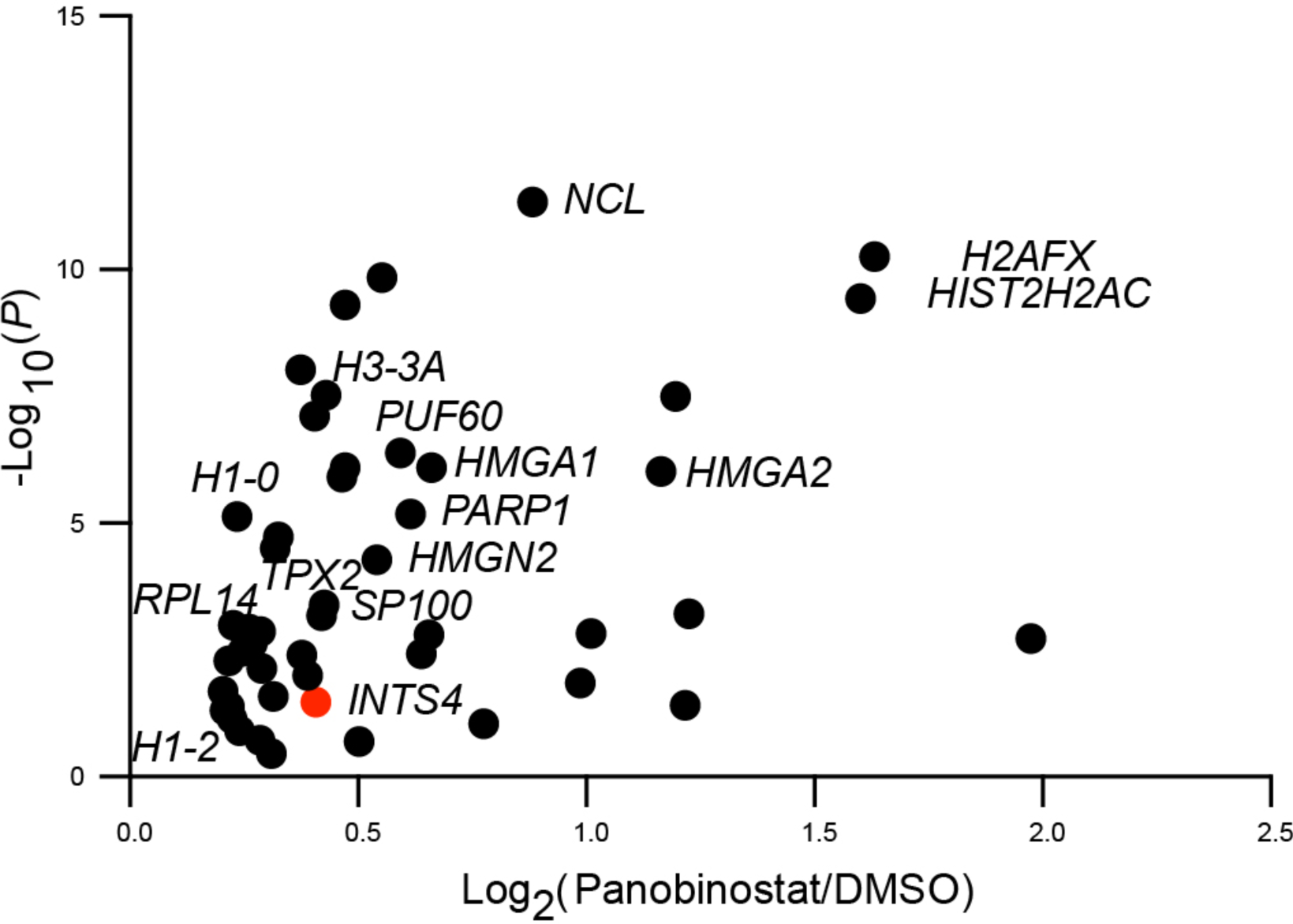
Acetylated targets enriched in HCT116 upon deacetylase inhibition. Enrichment plot for proteins with increased acetylation (Log_2_Fc> 0.4) in nuclei of HCT116 cells treated with panobinostat for 4 h with respect to DMSO control, derived from immune-enriched acetylated lysine containing peptides. Data represent hits from a single mass spectrometric run with nine biological replicates.

**Figure S12:**
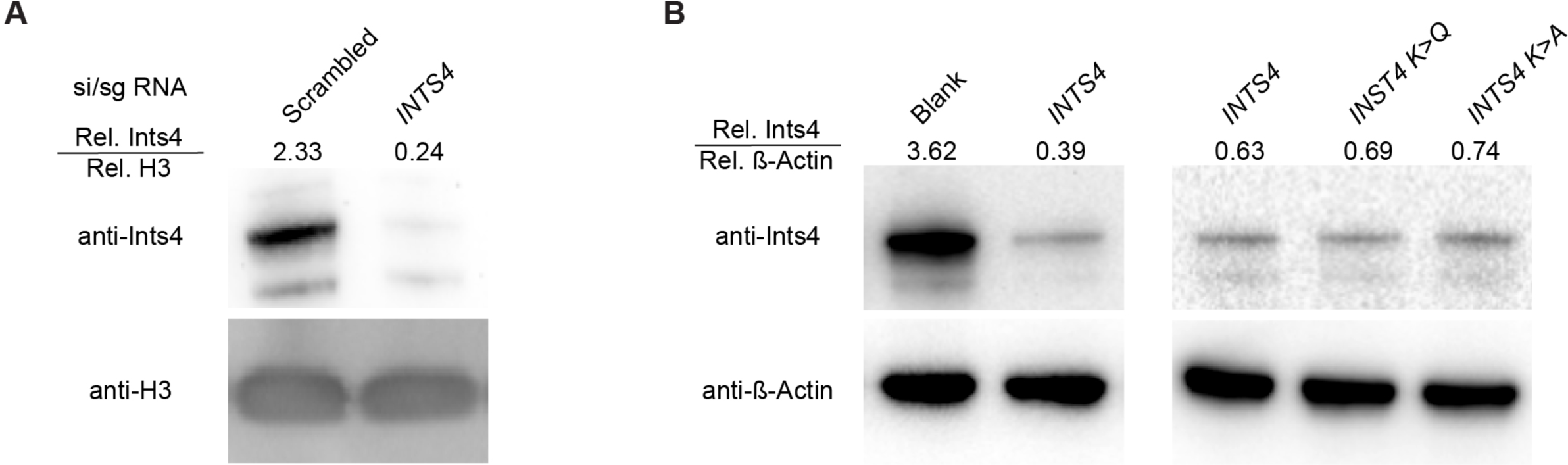
*INTS4* expression levels in knockdown and over expressing mutants. Western blot and quantitation for (**A**) *INTS4* knock-down in wild-type cells or (**B**) clones with over expression of ectopic wild-type *INTS4*, acetylation mimic (*INTS4 K>Q*) and acetylation null (*INTS4 K>A*) gene.

**Figure S13:**
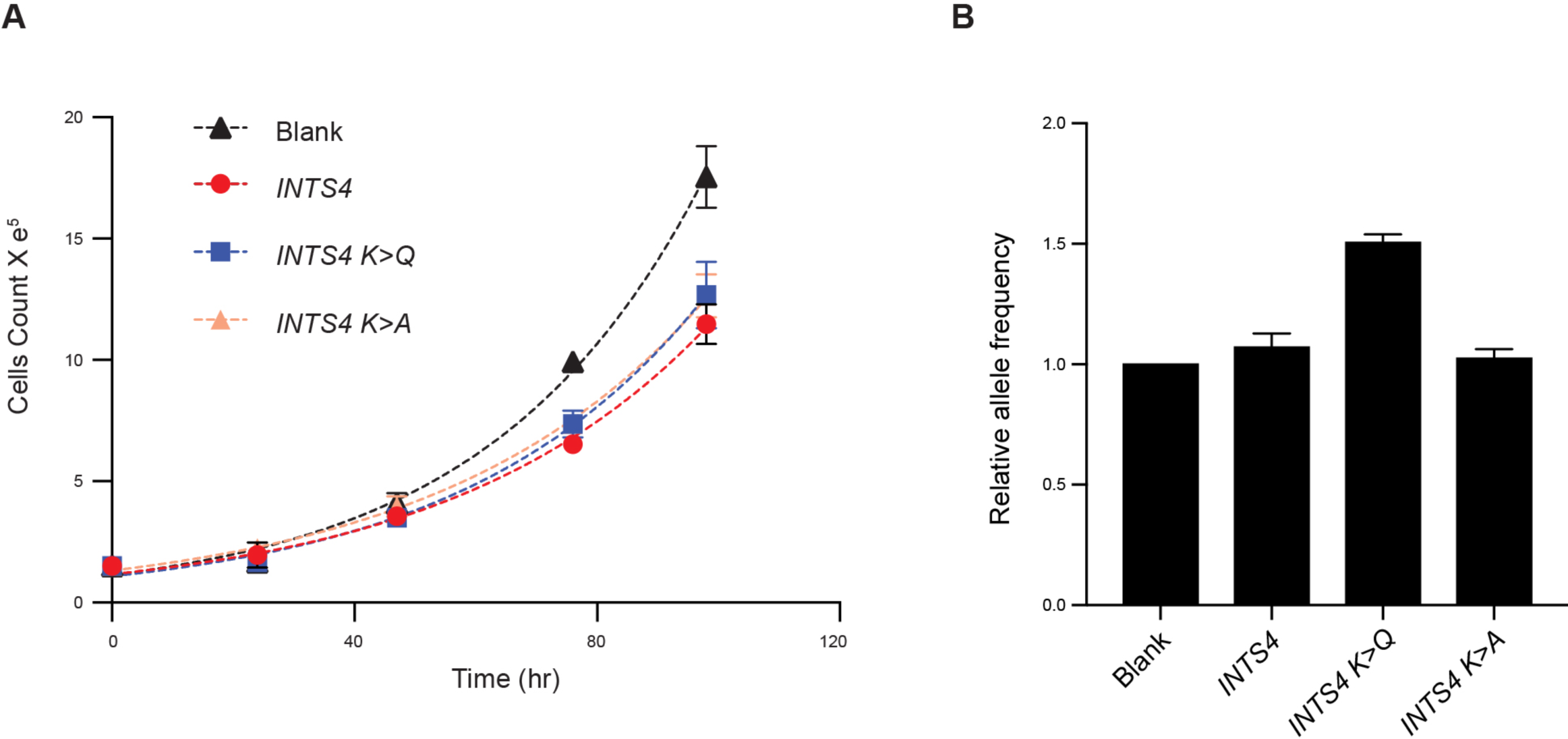
Effect of *INTS4* overexpressing on growth and allele frequency. **A**. Growth curve of monoclonal cell lines with no ectopic *INTS4* (Blank) or ectopic expression of either wild-type *INTS4*, acetylation mimic (*INTS4 K>Q*) and acetylation null (*INTS4 K>A*) mutant. The values represent mean ± SEM from 3 biological replicates. **B**. The normalized allele frequency in wild-type and mutant *INTS4* overexpressing clones, relative to Blank. The values represent mean ± SEM of relative allele frequency form DNA-FISH targeting multiple loci (Table S6).

Table S1: List of compounds in the small molecule library, related to figure S2.

Table S2: *P* values for changes in *ERRFI1* transcription site frequency with all the compounds in the library, related to figure 1D.

Table S3: Relative change in transcription site frequency and list of significant hits across three cell lines, related to figure 2A.

Table S4: Relative change in transcription site intensity and list of significant hits across three cell lines, related to figure 3A.

Table S5: List of hits from mass spectrometry, related to figure 6A and S11.

Table S6: Oligo information for DNA- and RNA-FISH probes, sg- and si-RNA and qPCR primers.

## Materials and Methods

### Cell culture

Human bronchial epithelial cells (HBECs), breast cancer cells MCF7 and colon HCT116 cells were grown in Keratinocyte-SFM medium (Gibco, 17005-042), MEM media (Gibco 11095-098) with 10% FBS (Gibco 10437-028) or MaCoy 5A (ATCC 30-2007) with 10% FBS, respectively. All cell culture media were supplemented with Penicillin/Strep (100units/mL Penicillin, 100micrograms/mL Streptomycin) and MEM media was additionally supplemented with 2mM glutamine (Gibco 25030-081).

For lentivirus production, HEK293FT cells were grown in DMEM (Thermo #11960-044) supplemented with 10% FBS (Thermo #10437), 1% penicillin-streptomycin and 2mM glutamine.

### Lentiviral construction and transfection

Lentiviral viral vector for dCas9-p300 (pHR_TRE3G-p300-dCas9-P2A-mCherry, Addgene: 138456), dCas9-VP64 (Addgene: 61422) and dCas9-Krab (Addgene: 85449) were purchased from the vendor. Lentiviral vectors for overexpression of wild-type and mutant *INTS4* were constructed as follows: *INTS4* was synthesized based on the cDNA sequence to code for a full length INTS4 protein (963 amino acid) with XbaI and NotI overhangs and cloned into the TOPO vector. Quick change was performed using Q5® Site-Directed Mutagenesis Kit (NEB: E0554S) to mutate the lysines at position 26-27 to glutamines or alanines. The wild-type and mutant *INTS4* coding sequences from TOPO vector were digested and purified for cloning between the XbaI and NotI sites in lentiviral expression vector pLV EF1alpha MCS IRES Cherry (Takara: 631987) using T4 DNA ligase (M0202S). Sequences of all vectors were confirmed by sanger sequencing.

HEK293FT cells (4X10^6^ in 10cm plate) were transfected with vectors containing the gene of interest (1.64pmol), pMD2g (0.72pmol), and pSPAX2 (1.3pmol) using 30ul of polyethylenimine (1mg/ml) as transfection agent. Viruses were collected and concentrated with Lenti-X Concentrator (Takara), flash frozen in liquid nitrogen, and stored in −80°C. Frozen aliquots of lentiviruses generated from HEK293 cells were quickly thawed in a 37°C water bath, and lentiviruses were applied to HBEC and removed after 24 h. Cells were propagated and ∼10^7^ cells were harvested for florescent activated cell sorting using a Sony MA900 Multiapplication cell sorter. Gates were set on cells with high signal from the florescent reporters: excitation of cells with dCas9-p300-2A-Cherry or INTS4-IRES-Cherry was done with a 561nm laser and emission assayed with a 617/30nm band pass filter, excitation for cells with dCas9-Krab-IRES-BFP or dCas9-VP64-2A-GFP, were done with a 405nm or 488nm laser, respectively, and emission assayed with a 525/50nm band pass filter. For single cell clones, about 500 sorted cells were propagated in a 15cm plate and colonies isolated using cloning cylinders for further characterization.

### Electroporation of synthetic guides and siRNA

2X10^5^ HBEC cells were washed with PBS and suspended in 10μl of buffer R (Thermo # NEON1) with 2.5μM synthetic guides for electroporation with Neon™ NxT Electroporation System using 4 pulses of 10ms at 1250V. For *INTS4* depletion 1μM of siRNA was additionally added in buffer R. Electroporated cells were diluted in 200μl of fresh SFM media for 10 min at room temperature before plating for experiments.

### RNA-FISH

10^4^ cells/well were plated in 384-well plate (Perkin Emler # 6057300) overnight and fixed for 15 min with 4% paraformaldehyde. Cells were washed twice for 5 min with 1X PBS before permeabilization with 70% ethanol for at least two hours at room temperature. Ethanol was removed with three rinses of 1X PBS. Cells were equilibrated with RNA-FISH wash buffer (10% Formamide, 2X SSC) for 30 min and then aspirated completely. Custom designed fluorescently labelled (Atto-647 or Quasar-670) RNA oligo (Biosearch Technologies) directed to the introns of target gene were diluted to 10nM in Hybridization buffer (10% Formamide, 2X SSC, 10% dextran sulfate) and incubated overnight with cells at 37°C. Excess probes were removed by two 10 min washes with RNA-FISH wash buffer (10% Formamide and 2X SSC) and one 5 min wash with 2X SSC (0.3 M Sodium Chloride and 0.03M Sodium Citrate, pH7). Cells were further washed with 1X PBS for 10 min and stained with DAPI. For high-throughput RNA-FISH across multiple 384-well plates, cell seeding, fixation and fluid transfers were done with either multichannel electronic pipettes and/or automated liquid handlers (Biotek EL406 and BlueCatBio).

### DNA-FISH

Cells were plated and fixed like the RNA-FISH protocol described above. Permeabilization was performed with 0.5% Saponin and 0.5% Triton X 100 in PBS for 20 min and detergent washed off with two PBS washes. Cells were next incubated with 0.1N HCL for 15 min and washed with 2X SSC for 5 min. Cells were further incubated in 50% formamide in 2XSSC for at least 1 hour. The formamide solution was aspirated and 2 μl of custom probes and 8 μl of Hyb buffer from Empire Genomics (https://empiregenomics.com/) added to each well. Denaturation was done at 85°C for 7 min and cells were shifted to 37°C for hybridization overnight. Excess probes were then washed off with 3 washes of 1X SSC (150mM sodium chloride and 15mM sodium citrate) and 3 washes of 0.1X SSC for 5 min each at 42°C. Finally, nuclei were stained with DAPI and imaged in PBS.

### RT-qPCR

HBEC cells (5e^5^) from different treatments were mixed mouse RPE1 cells (5e^5^) for harvesting total RNA using RNeasy kits (Qiagen #74104) for reverse transcription with iScriptTM reverse transcription supermix (Bio-Rad #1708840) and subsequent qPCR with iTaq Supermix (Bio-Rad #1725120). The relative abundances from a standard plot were normalized to mouse actin between samples.

### Small molecule library screen

The library (Cayman # 11076) was diluted manually to a working stock of 40μM in 1X PBS and aliquoted in a donor plate (Echo 384PP 2.0, Bio Lab). The inhibitors were transferred to the designated wells in 384-well plates with 80% confluent cells using a ECHO525 Acoustic Liquid Handler (Beckman # 001-10080) to a final concentration of 1μM. The cells were fixed after 4 h of inhibition and processed for nscRNA-FISH as describe above.

### High-throughput image acquisition

For live cell bursting assay, images were acquired using an automated high-throughput dual spinning disk microscope (Yokogawa Cell Voyager 7000S) in a 37°C, 5% CO2 and 80% humidity environment. Excitation was done with a 488 nm excitation laser and a quad-band dichroic mirror (405/488/561/604 nm). Images were acquired by 60X water immersion objective (1.2 NA) through a 525/50 nm bandpass emission filter and recorded on 16-bit Andor Neo 5.5 sCMOS camera with 2 × 2-pixel binning, resulting in a XY pixel size of 216.6 nm. A Z-stack of 10 images with 0.5 μm step size was collected for each field of view (FOV). Time-lapse imaging at 100 s intervals was done for up to 12 h. Typically, around 20-28 FOV were acquired per experiment. Flat-field correction and maximum intensity projections were performed on-the-fly by the Yokogawa controlling software and a 2D time-lapse (2D-t) image sequence were generated per FOV.

For imaging nscRNA-FISH, the acquisition paraments were similar to the live-cell assay described above with the following differences. Imaging was done at room temperature, fluorophore excitation of DAPI nuclear stain and Atto-647/Quasar-670 dye in RNA-FISH probes was done with 405nm and 640nm laser, respectively, and signals were sequentially acquired with 445/45nm and 679/29nm band pass filters, respectively, for sufficient FOVs to image 500-1000 cell/well.

### Image analysis

For nscRNA-FISH, images were analyzed using Columbus 2.8.2 (PerkinEmler). Automated nuclear segmentation was done on the maximum intensity projection images from the DAPI channel using the following filtration criteria: Common threshold: 0.4; area: >30μm^2^; splitting coefficient: 7; individual threshold: 0.4; contrast: > 0.1. Partial cell images at the edge of the FOV were removed from analysis. The final regions assigned as nuclei in the preceding steps were subjected to automated spot segmentation on the maximum intensity projection images from the far-red channel (679/29nm) using the following selection criteria: radius: <6.9 pixel; contrast: >0.24; uncorrected spot to region intensity: 1.5-0.2.5 (dependent on the gene); distance: >11.8 pixel; spot peak radius: 3 pixel. The stats from the batch analysis of the entire screen were exported to get cell-level data for number and intensity of TS. For intensity analysis we focused on the relative change in-lieu of absolute intensity as it both normalizes for the differences in the number of probes between genes and variations in FISH efficiencies across experiments. Custom scripts in the R software were used to extract and process imaging data for figure preparation. Any RNA-FISH artifacts arising from debris that falsely segment as high intensity spots were filtered out post imaging by filtering the intensity values between 1-99% of the distribution for each well and the distribution further checked for anomalies. The *P* values for all treatments were normalized with the Benjamini-Hochberg procedure to reduce false discovery rate^86^.

Image analysis of live-cell bursting was done as described previously^16^. Briefly, the image processing workflow consisted of for major modules: (i) segmentation^87, 88^ and two-dimensional tracking^89^ of individual HBEC nuclei in time-lapse FOVs; (ii) automatic rigid-body registration of individual tracked nuclei^90^; (iii) automatic detection (GDSC-SMLM ImageJ Plugin, University of Sussex, https://sites.imagej.net/GDSC-SMLM/) and tracking of active TS in individual living cells^89^; and (iv) extraction of TS intensity trajectories over time. Deep-learning algorithms were incorporated to improve both the cell and spot segmentation. All transcription-site trajectories were manually verified and filtered using an interactive platform to ensure any spurious traces are omitted for kinetic analysis. All workflows were implemented using the open-source workflow orchestration and management software, Konstanz Information Mining (KNIME, 64-bit, Version 3.2.1)^91^ with KNIME Image Processing nodes (KNIP, Version 1.5.3.201611190650)^92^, R (64-bit, Version 3.3.1) and Python (64-bit, Version 2.7.12) scripting KNIME nodes.

### Hidden Markov model analysis

To extract the On- and Off- times from the live cell intensity traces, each trace was first normalized to a peak height of one. Next, normalized data were fed to the IDL program ‘HMM_V5_4_2_text.sav’ described in^93^ (https://sites.psu.edu/smbiophysics/hmm-code-and-manual/). The program uses Hidden Markov Model (HMM) algorithm for fitting two states and outputs a binary trace representing the times when the gene is active and inactive. Batch analysis of multiple traces for each condition was used to generate the normalized distribution of On- and Off- times of bursting.

### Cut and Run

The CUT and RUN protocol was adapted from Mura and Chen^94^. Briefly, 1.8X10^5^ cells were plated overnight in a standard 96-well plate for gentle fixation with 0.5% PFA for 2min, followed by neutralized with 125mM glycine for 2 min and washed three times with PBS. Cell were permeabilized with permeabilization buffer (0.1% Triton X-100, 20 mM HEPES-KOH pH 7.5, 150 mM NaCl, 0.5 mM Spermidine, 0.01% Digitonin) containing proteinase inhibitor (Roche, 04 693 132 001) for 15 min at room temperature and washed once with permeabilization buffer. Cells were incubated in 100μl of permeabilization buffer with 2μg of anti-H3K27ac (Abcam # ab4729) for at least 2 h at 4°C. The unbound antibodies were washed away twice with 100μl of cold permeabilization buffer for 3 min before adding 1.5μl of pAGMnase (Signaling Technology, 86652) in 100μl of permeabilization buffer at room temperature. Unbound MNase was removed with two 5 min washes with the permeabilization buffer. During the second wash of this stage, the plate was placed over wet ice at 4°C. Cuts surrounding the antibody bound sites were induced by activating pAG-MNase with 150µl ice-cold permeabilization buffer containing 5 mM CaCl2 at 4°C for 30-40 min and digestion was halted by adding 50µl 4× STOP solution (680 mM NaCl, 40 mM EDTA, 8mM EGTA, 100µg/ml RNase A and 0.1% Triton X-100) containing 2.5ng of spike-in DNA (Cell Signaling Technology, 86652). The plate was incubated at 37°C for 10 min to facilitate the release of digested DNA fragments from the nucleus. The supernatant was transferred to DNA LoBind® Tubes (Eppendorf #22431005). Proteinase K (40ug) and SDS (final 1%) was added to the supernatant and incubated at 55°C for 2 h followed by 65°C for 7 h, for un-crosslinking. The proteins were precipitated using 300µl of phenol/chloroform and phenol was then removed with another chloroform wash. MNase digested fragments were precipitated with 500µl of ethanol in presence of 0.3M sodium acetate and 20µg of glycogen. To increase yield, samples were incubated at −80°C for 2 h before spinning at 20K rcf for 1.5 h at 4°C. The precipitated DNA was washed once with 70% ethanol and suspended in 30μl 0.1 X TE (1mM Tris-HCl and 0.1mM EDTA) for library preparation with NEBNext® Ultra™ II DNA Library Prep Kit for Illumina® (NEB # E7645S) following the manufacturer’s protocol. The final library was suspended in 33μl of 0.1X TE and quality checked by running 1μl on a high sensitivity Agilent D1000 Tape Station with a prominent mono- and a small di-nucleosomal peak and further diluted in 150 μl of distilled water for qPCR.

### Spike-in ChIP-Seq

ChIP-seq was performed as previously described^95, 96^. In brief, 10^7^cells were fixed on plate with 1% paraformaldehyde for 15 min and neutralized with 125mM glycine. Fixed cells were washed twice with ice-cold PBS and scraped off in ice-cold PBS containing Protease Inhibitor Cocktail (Sigma–Aldrich #P2714). Cells were pelleted and then resuspended in ChIP Lysis Buffer (0.5% (w/v) sodium dodecyl sulfate (SDS), 8 mM EDTA, 40 mM Tris-HCl (pH 8), protease inhibitor cocktail). Cells were kept on ice for one hour before sonication at 4°C (Bioruptor, Diagenode) to an average DNA length of 200–500 bp by 15 pulses of 10 sec on and 10 sec off at high intensity. Cell debris was removed by centrifugation at top speed for 20 min and chromatin concentration recorded using OD_260_. Samples were diluted to a final concentration of 200μg/ml in ChIP dilution buffer (0.01% (w/v) SDS, 1.2 mM EDTA, 16.7 mM Tris-HCl (pH 8), 1.1% (v/v) Triton X-100, 167 mM NaCl, protease inhibitor cocktail) and 600μg chromatin was used for each immunoprecipitation. 4 μg of anti-H3K27Ac coupled to 50 μl Pierce™ Protein A/G Magnetic Beads (Thermo #88802) was added to each sample and rotated overnight at 4°C. For spike-in normalization 4μg of drosophila chromatin (Active motif # 53083) and 2μl spike-in antibody (Active motif # 61686) were added per sample. The beads were harvested by magnets and washed once for 10 min in the cold room by rotation with 1 ml of ChIP Low Salt Wash Buffer (0.01% (w/v) SDS, 2 mM EDTA, 20 mM Tris-HCl (pH 8), 1% (v/v) Triton X-100, 150 mM NaCl), 1 ml of ChIP High Salt Wash Buffer (0.01% (w/v) SDS, 2 mM EDTA, 20 mM Tris-HCl (pH 8), 1% (v/v) Triton X-100, 500 mM NaCl), and 1 ml of ChIP LiCl Wash Buffer (250 mM LiCl, 1% (w/v) IGEPAL, 1% (w/v) sodium deoxycholate, 10 mM Tris-HCl (pH 8), 1 mM EDTA). Finally, the beads were washed twice for 2 min in the cold room by rotation with 1 ml of TE buffer (1 mM EDTA, 10 mM Tris-HCl (pH 8)). Antibody-bound chromatin fragments were digested and crosslinks reversed with Reversal Buffer (0.0075% (w/v) SDS, 200 mM NaCl, 10 mM EDTA, 50 mM Tris-HCl (pH 7.5), 25µg proteinase K (Thermo #E00491) by incubating at 50°C for two hours with shaking and subsequently seven hours at 65°C. ChIP DNA samples were purified with phenol-chloroform extraction and ethanol precipitation with glycogen (Thermo #R0551) as carrier. ChIP-seq libraries were generated using NEBNext® Ultra™ II DNA Library Prep Kit for Illumina® (NEB # E7645S) following the manufacturer’s protocol. Libraries were multiplexed for single-end sequencing using the NextSeq500 High Output Kit v2 (FC-4042005, 75 cycles) on an Illumina NextSeq500.

### Analysis of deep sequencing

Data was process as described previously^97, 98^. Briefly, single-end reads were aligned to the genome sequence of concatenated human (hg38) and spike-in *Drosophila* (dm6) genomes using Bowtie 2^99^ with specified options “–no-mixed” and “–no-discordant”. PCR duplicates and reads mapped more than once were removed with SAMTools^100^. The mapped hg38 reads were down sampled by multiplying the ratio of minimum dm6 reads between all samples to the mapped dm6 reads for respective samples. The maximum value for the down-sampling factor thus equals 1 for sample with lowest dm6 reads and its hg38 count remains unchanged. Read counts for remaining samples were down-sampled based on the ratio described above. Raw and processed files are available on GEO data base (accession number GSE254866). Read coverage and reproducibility across replicates was analyzed using multiBamSummary and plotCorrelation functions from deepTools suite^101^. Genome coverage tracks were generated using the pileup function from MACS2^102^. The meta-gene plot coordinates were generated using the computeMatrix and plotProfile/plotHeatmap functions from deepTools and plotted with custom R-scripts.

### Mass Spectrometric analysis

Nuclei were isolated as described previously^103^. Briefly, confluent cells in a 15 cm dish were treated with either 1μM Panobinostat or DMSO, then harvested by trypsinizing and washing twice with cold PBS. About 5X10^6^ cells were suspended in cold hypotonic lysis buffer (HLB; 10 mM Tris (pH 7.5), 10 mM NaCl, 3 mM MgCl2, 1% (vol/vol) NP-40 and 10% (vol/vol) glycerol) with 1X Thermo Scientific™ Halt™ Protease and Phosphatase Inhibitor Cocktail (Thermo #78443) and incubated on ice for 15 min. The cells were vortexed for 20 sec and again incubated on ice for 10 min. Nuclei were spun at 10^3^ rcf for 3 min at 4°C and suspended in HLB. The cells were washed twice with HLB and frozen for transportation. A small number of nuclei were removed for staining with DAPI and ER-Tracker™ Red (Thermo # E34250) to check for loss of ER staining.

The nuclear lysates were precipitated by addition of TCA (4:1 v/v ratio of TCA/sample). The protein pellet was resuspended in 700µL of EasyPep lysis buffer (Thermo # A45735) containing 1X deacetylase inhibitors (Active Motif #37494) and treated with TCEP and chloroacetamide to a final concentration of 16mM and 32mM, respectively. The reaction was incubated at 25°C for 1 h in the dark and incubated with 7.5µg of trypsin/LysC (Thermo # 90051) at 37°C for 16 h. TMTpro labels (500µg; Thermo # A52045) were added to each sample and incubated for 1 h at 25°C with shaking. Excess TMTpro was quenched with 50μL of 5% hydroxylamine and 20% formic acid for 10 min and samples were then combined and cleaned using three EasyPep Maxi (Thermo# A45734) columns as described in the manual. Peptides were eluted in 3ml of elution buffer provided in the kit for each column then combined and 25µL was dried and saved for total protein analysis. The rest was split into 8 aliquots dried and enriched for acetylated peptides using the PTMScan® HS Acetyl-Lysine Motif (Ac-K) Kit (Cell Signaling Technology #46784) as described in the manual and peptides were eluted twice with 200µL of 50% acetonitrile (ACN), 0.5% trifluoroacetic acid (TFA) and eluted fractions were combined and dried.

Total and acetylated peptides were analyzed in triplicate using Dionex U3000 RSLC in front of a Orbitrap Eclipse (Thermo) equipped with an EasySpray ion source. Solvent A consisted of 0.1% formic acid (FA) in water and Solvent B consisted of 0.1% FA in 80% ACN. The loading pump consisted of Solvent A and was operated at 7 μL/min for the first 6 min of the run then dropped to 2 μL/min when the valve was switched to bring the trap column (Acclaim™ PepMap™ 100 C18 HPLC Column, 3μm, 75μm I.D., 2cm, PN 164535) in-line with the analytical column EasySpray C18 HPLC Column, 2μm, 75μm I.D., 25cm, PN ES902). The gradient pump was operated at a flow rate of 300nL/min. For all samples, a linear LC gradient of 5-7% B for 1 min, 7-30% B for 133 min, 30- 50% B for 35 min, 50-95% B for 4 min, holding at 95% B for 7 min, then re-equilibration of analytical column at 5% B for 17 min. The MS methods employed the TopSpeed method with three FAIMS compensation voltages (CVs) and a 1 second cycle time for each CV (3 second cycle time total) that consisted of the following: Spray voltage was 2200V and ion transfer temperature of 300°C. MS1 scans were acquired in the Orbitrap with resolution of 120,000, AGC of 4e^5^ ions, and max injection time of 50 ms, mass range of 350-1600 m/z; MS2 scans were acquired in the Orbitrap using TurboTMT method with resolution of 15,000, AGC of 1.25e^5^, max injection time of 22ms, HCD energy of 38%, isolation width of 0.4Da, intensity threshold of 2.5e^4^ and charges 2-6 for MS2 selection. Advanced Peak Determination, Monoisotopic Precursor selection (MIPS), and EASY-IC for internal calibration were enabled and dynamic exclusion was set to a count of 1 for 15 sec. The only difference in the methods was the CVs used, the first used CVs of −45, −60, −75, the second use CVs of −50, −65, −80, and the third used CVs of −55, −70, −85.

All MS files were searched with Proteome Discoverer 2.4 using the Sequest node. Data was searched against the Uniprot Human database from Feb 2020 using a full tryptic digest, 2 max missed cleavages, minimum peptide length of 6 amino acids and maximum peptide length of 40 amino acids, an MS1 mass tolerance of 10 ppm, MS2 mass tolerance of 0.02 Da, variable oxidation on methionine (+15.995 Da), TMTpro (+304.207) on lysine and peptide N-terminus and acetylation on lysine (+42.011 Da) and fixed carbamidomethyl on cysteine (+57.021). Percolator was used for FDR analysis and TMTpro reporter ions were quantified using the Reporter Ion Quantifier node and normalized on total peptide intensity of each channel. The raw intensity data from mass spectrometry is available at ftp://MSV000094611@massive.ucsd.edu.

To filter hits with increased acetylation upon deacetylase inhibition, first, the acetyl-peptides intensities arising from each protein were summed for each replicate of DMSO and panobinostat treated cells. Next, the mean intensity between replicates of panobinostat and DMSO treated cells was obtained and a cutoff > 0.2 for the Log_2_ fold change relative to DMSO control was used to filter enriched hits. For HBECs, the experiment was performed twice with 5 and 9 replicates. Enriched hits that were shared between both experiments and had robust signal for both DMSO and panobinostat treated replicates were used for calling high confidence hits. For HCT116 cells the experiment was performed once with 9 replicates of DMSO and panobinostat treated cells and hits filtered with a cutoff > 0.4 for the Log_2_ fold change relative to DMSO.

## Statistical analysis

Custom R-scripts and software GraphPad Prism was used for analyzing data. The number of replicates and variability parameters can be found in the corresponding figure legends.

## Data and code availability

Raw and processed files are available for download from NCBI GEO, accession number GSE254866 The image analysis pipelines in the software Knime and Columbus along with the associated R-scripts used for statistical analysis are available on https://github.com/CBIIT/MIstelilab_Sood-et-al.

